# Photosynthetic protein classification using genome neighborhood-based machine learning feature

**DOI:** 10.1101/2020.01.09.898809

**Authors:** Apiwat Sangphukieo, Teeraphan Laomettachit, Marasri Ruengjitchatchawalya

**Affiliations:** Bioinformatics and Systems Biology Program, School of Bioresources and Technology, King Mongkut’s University of Technology Thonburi (KMUTT), Bang Khun Thian, Bangkok 10150, Thailand; Biotechnology program, School of Bioresources and Technology, KMUTT, Bang Khun Thian, Bangkok 10150, Thailand; School of Information Technology, KMUTT, Bang Mod, Thung Khru, Bangkok 10140, Thailand

**Keywords:** Photosynthesis, Genome neighborhood, Machine learning, Protein function

## Abstract

Identification of novel photosynthetic proteins is important for understanding and improving photosynthetic efficiency. Synergistically, genomic context such as genome neighborhood can provide additional useful information to identify the photosynthetic proteins. We, therefore, expected that applying the computational approach, particularly machine learning (ML) with the genome neighborhood-based feature should facilitate the photosynthetic function assignment. Our results revealed a functional relationship between photosynthetic genes and their genomic neighbors, indicating the possibility to assign functions from their genome neighborhood profile. Therefore, we created a new method for extracting the patterns based on genome neighborhood network (GNN) and applied for the photosynthetic protein classification using ML algorithms. Random forest (RF) classifier using genome neighborhood-based features achieved the highest accuracy up to 94% in the classification of photosynthetic proteins and also showed better performance (Mathew’s correlation coefficient = 0.852) than other available tools including the sequence similarity search (0.497) and ML-based method (0.512). Furthermore, we demonstrated the ability of our model to identify novel photosynthetic proteins comparing to the other methods. Our classifier is available at http://bicep.kmutt.ac.th/photomod_standalone, https://bit.ly/2S0I2Ox and DockerHub: https://hub.docker.com/r/asangphukieo/photomod

## Introduction

Photosynthesis is a multistep process comprising solar energy harvest, excitation energy transfer, energy conversion, electron transport, in particular, the transport of electrons flow from water to NADP+ and other electrochemical energy via photosystems and a series of enzymatic reactions required to synthesize components for cellular metabolism and environmental adaptation. Recently, photosynthetic prokaryotes, in comparison with plants, have attracted considerable attention for a wide range of renewable energy, agricultural and environmental applications aiming for sustainable development (Pathak, et al., 2018), probably due to their excellent productivity. However, it has been reported that the photosynthetic conversion efficiency falls around 6% of the total incident light. Maximizing photosynthetic efficiency is a major challenge in the current efforts to manage and/or engineer the photosynthetic organisms (Work, et al., 2012). To achieve this goal, all photosynthetic components and their roles in photosynthesis need to be clarified beforehand.

Basically, photosynthesis-related genes can be identified by two mains approaches: experimental and computational approaches. Although experimental approaches have successfully identified many components (Wegener, et al., 2008), time and cost wastages are the major limitations. Besides, many photosynthetic components are temporarily present (Eaton-Rye and Sobotka, 2017), and even the deletion of those genes might have no effect on photosynthetic growth (Wegener, et al., 2008). These limitations push forward the development of various computational approaches. These approaches have been used to narrow down and specify the gene function before the validation by expensive experimental approaches. Sequence similarity search is conventionally used for gene annotation, and its validity depends on prior knowledge. By searching against known protein databases, (i.e. NCBI databases), proteins involved in photosynthesis are hypothetically annotated with unknown functions and many times misidentified (Nagashima and Nagashima, 2013). It has been shown that 70% of the close homologs of photosynthetic proteins come from non-photosynthetic organisms (Ashkenazi, et al., 2012) resulting in unsatisfied results in the attempt of photosynthetic protein identification. Unlike the conventional approach, ML provides insight into the functional classification of novel proteins with no homology to proteins of known function (Han, et al., 2005). Various ML methods have been developed with different learning features and algorithms. SCMPSP (Vasylenko, et al., 2015) calculates physicochemical property from dipeptide and amino acid composition optimized by genetic algorithm for characterizing photosynthetic proteins. SVMprot (Li, et al., 2016) uses support vector machine (SVM) to learn the physicochemical features of representative proteins to predict protein function including photosynthesis function. Recently, DeepGO (Kulmanov, et al., 2018) uses deep neuron networks to predict protein functions from protein sequences and protein-protein interaction networks. However, poor prediction performance, especially in relation to biological process (Skunca, et al., 2012), has necessitated the use of additional information such as gene cluster to infer protein function (Zheng, et al., 2011).

Analysis of prokaryote genomes has revealed that genes, including those involved in the photosynthesis, tend to form clusters (Nagashima and Nagashima, 2013; Rogozin, et al., 2002). It has been suggested that genes located in the same cluster participate in the same biochemical network (Bergeron, et al., 2008) or exhibit the same expression pattern (Semon and Duret, 2006). Thus, analysis of such conserved clusters provides useful information for gene annotation and genome evolution. Several methods have been developed to uncover conserved gene clusters and identify the functional connection in such group of genes (Lemay, et al., 2012; Rogozin, et al., 2002; Zhao, et al., 2014). The general approach starts by identifying an orthologous set of a given gene in all available genomes. Next, the neighboring genes are extracted and ranked by the frequency of their occurrence. The neighboring genes are expected to have a functional connection to the orthologs if there is a statistically significant association between them (Galperin and Koonin, 2014). Instead of using the number of occurrences, a tree-based probabilistic method was developed and showed better performance (Zheng, et al., 2005). However, the difficulty in handling large scale data limits the application of this genomic context. Lately, GNN was shown to cope with large scale prediction of enzymatic activities of uncharacterized enzymes (Zhao, et al., 2014). However, the use of this method requires prior knowledge of the protein families and complicated interpretation (Zallot, et al., 2016).

Therefore, we expected that applying ML approach with genome neighborhood-based feature should facilitate the photosynthetic function assignment. In the first section of this study, we observed functional relationships between photosynthetic genes and their neighbors to indicate the possibility of using information from neighboring genes to infer the photosynthetic function. Next, gene neighborhood patterns were extracted by our newly developed method based on GNN. The patterns were applied as features for learning by ML algorithms. Then, the new model was fine-tuned until it reached the highest performance. The application of the model was demonstrated by comparing the prediction performance with other available tools and also, by predicting 12 novel photosynthetic proteins, which are recently identified by experiments.

## Material and Methods

### Gene neighborhood calling

As described in Text S1, we retrieved 154 completed photosynthetic prokaryote genomes from the NCBI database. Genes in each genome were identified and an in-house python script was used for the selection of neighboring genes in each genome. Genes on the same strand were considered neighbors if they are within an intergenic distance of not more than 250 bp (Lathe, et al., 2000) or are overlapping (Figure 1A). In addition, when the gap between the first genes of two neighborhood gene clusters with divergent directions is within a range of 200-1000 bp, the two clusters were merged into the same neighborhood gene cluster, based on the operon interaction concept. It was shown that pairs of operons that regulate each other and/or those that are co-regulated tend to lie in the range of 200-1000 bp (Warren and ten Wolde, 2004).

**Figure 1.**
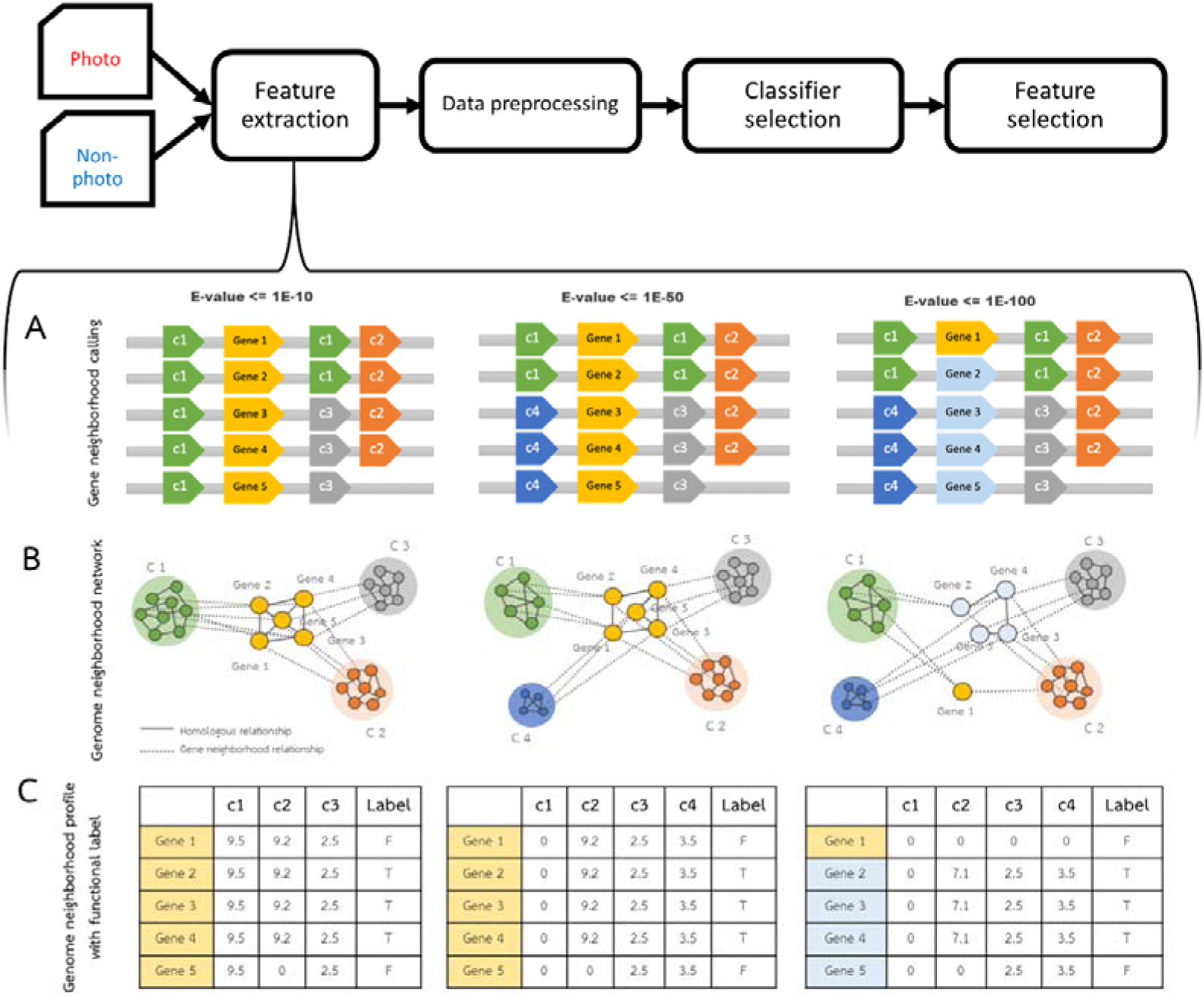
Protocol for building the model of photosynthetic protein classification and the demonstration of the feature extraction method. The protocol consists of dataset building, feature extraction, data preprocessing, classifier selection and feature selection. (A) Genome neighborhoods are called by using intergenic distance criteria in the feature extraction step. Genes in the same homologous group are indicated by the same color label. The query genes conserved in different five genomes are labeled by Genes 1-5. (B) Relationships between query genes and their neighbors are displayed in GNN. Straight-line represents a homologous relationship, while the dashed line represents genome neighborhood relationships. Color labeled on each node represents a homologous group corresponding to gene color in (A). (C) The genome neighborhood profile of each query gene is represented in table format. The value in the table is the Phylo score, which represents the level of gene neighborhood conservation. E-values (1E-10, 1E-50 and 1E-100) are the thresholds for family classification of proteins coding from the genes.

### Genome neighborhood network

Sequence similarity network (SSN) was applied to discriminate the protein function (Atkinson, et al., 2009) by using different stringent E-value thresholds to construct the group-wise sequence similarity relationships. This approach enables functional and evolutional analysis of large protein datasets and is also applied to genome neighborhoods to guide the discovery of novel metabolic pathways (Zhao, et al., 2014). In this work, three different E-value cutoffs of Blastp were employed (1E-10, 1E-50 and 1E-100) to all proteins from the 154 genomes. Then the Blastp outputs were used as input in Markov cluster algorithm (MCL) to cluster proteins into families (see Text S1). Figure 1B visualizes genome neighborhood networks, where homologous relationships from the SSN at different E-value cutoffs were integrated with gene neighborhood relationships from the gene neighborhood calling step.

### Gene neighborhood profile generation

Figure 1C shows the gene neighborhood profile scores (calculated as Phylo scores) of each conserved neighbor cluster IDs with different E-value cutoffs. Note that the Phylo score of neighboring genes was calculated if the query genes (e.g., genes 1-5 in Figure 1) and the neighboring genes are conserved in at least three genomes. The calculation of Phylo scores includes three main steps: i) performing bidirectional Blast best hit of all pairwise genomes, ii) generating a pairwise metric of a proportion of shared gene content and iii) constructing a phylogenetic tree using genomes containing conserved gene neighbors. Figure S1 outlines the steps to calculate the Phylo scores. Here we describe each step in detail.

Pairwise comparisons of all 154 proteomes (Table S1) were performed using Blastp and the shared gene content of each pair of genomes was identified by reciprocal best hit, implemented by an in-house python script. The pairwise distance between genomes was calculated by *d = -ln(s)*, where *s* is the proportion of shared gene content of the two genomes divided by the average number of proteins between the two genomes. Then, the phylogenetic tree of the genomes, in which the gene neighbors were identified, was constructed using the pairwise distance profiles that were created, i.e. genome distance matrix. Biopython package in Python (Bio.Phylo.TreeConstruction) was used to generate the distance matrix and the phylogenetic tree. The phylogenetic tree was built by the neighbor-joining algorithm. The Phylo score was determined by the summation of the total branch length of the tree.

When the stringency was increased from 1E-10 to 1E-50, the neighboring genes of the query genes (Genes 1-5) split from three to four clusters as shown by GNN (Figure 1B). As a result, no Phylo score was assigned to c1 because the Phylo score was calculated (neighboring genes are considered conserved) only if genes are found as neighbors in at least three genomes. In addition, a new cluster (c4) that was split from c1 was added with a corresponding Phylo score. When the E-value was increased to 1E-100, the cluster of the query separated into two clusters (yellow and light blue clusters). Gene 1 now became the only member of the yellow cluster and as a result, no Phylo score was assigned. Accordingly, the Phylo score of the gene cluster c2 of the light blue query decreased because the number of genomes that light blue cluster genes and c2 cluster genes are conserved decreased from four to three. The last column in the table is the label of the protein function (T= photosynthesis, F= non-photosynthesis). As demonstrated, the low stringent E-value yields similar patterns of neighboring genes, thus it is impossible to separate the protein function. A significant difference in the patterns was observed in more stringent criteria. Thus, it was expected that when the three tables are combined together, they can separate the function of the query proteins. The point of this example is to show how the patterns of neighboring genes separate the true gene function from the background noise, even if the query sequences are closely related.

Practically, the Phylo score is numeric and diverse depending on the distance of the genomes, thus, to make it easy to understand by the machine, the normalization method was applied. Quartile was carried out to discretize all Phylo scores (from the entire neighboring gene of photosynthetic genes) to four levels. Phylo scores in Level 0 (Phylo score = 0), level 1 (first quartile), level 2 (second quartile) and level 3 (third quartile) indicate no gene conservation, low gene conservation, intermediate gene conservation, and strong gene conservation, respectively. After the process was computed, the Quartile 1 and Quartile 3 cutoff points of 0.41079 and 2.61799 respectively were found. The normalized Phylo score tables from different E-value criteria were combined and used to build models.

### GNN-based model construction for photosynthetic protein classification

#### Photosynthetic and non-photosynthetic protein dataset building

The first step of the approach involves generating input data. The photosynthetic proteins (6,430 sequences) and non-photosynthetic proteins (6,430 sequences) were retrieved from UniprotKB, which is a database of manually annotated protein sequences. The protocol was shown in Text S1 and Text S2.

#### GNN feature extraction and data preprocessing

The gene neighborhood profile was created for each protein sequence in the dataset as described in the method section. Briefly, the protocol contains three main steps as 1) calling gene neighbors, 2) measuring Phylo score and 3) normalizing the score. It was found that our dataset was largely redundant after calling gene neighborhood features. Therefore, data preprocessing was carried out by removing duplicated data, resulting in 1,176 unique instances (318 photosynthetic instances and 858 non-photosynthetic instances) of the total 12,858 instances. A total number of 18,849 features, cluster IDs, was obtained. This dataset represents two common problems encountered in real-world classification namely class imbalance and high-dimensional data. We carefully considered these problems by choosing the evaluation metric (Text S3) and classifier that are able to attenuate these problems. The dataset was separated into two parts: 1) 90% of the data for model construction and 2) 10% of the data as an independent test set.

#### Classifier selection

Different classifiers were included in this study according to their prediction performance on high-dimensional data (Amancio, et al., 2014; Caruana and Niculescu-Mizil, 2006). The first, a probabilistic-based classifier called Bayesian Network (BN) (Friedman, et al., 1997; Molchanov, et al.), of which structure is in the form of a directed acyclic graph, which allows attribute-attribute interaction, thereby improving classification performance, especially for large datasets. The second is a support vector machine (SVM) classifier named Sequential Minimal Optimization (SMO). The SVM-based classifier is known to handle high-dimensional data, such as the microarray classification, well (Nanni, et al., 2012). The last algorithm is a decision tree-based classifier named random forest (RF) (Breiman, 2001), which uses the vote from classifier multiple.

#### Feature selection

The last step of the model development involved model optimization, by which the optimal feature subsets were determined in order to make the model more robust and reduce the chance of fitting noise in the learning algorithm. Two different schemas were applied to the best-performed classifier selected from a previous step (Classifier selection). Firstly, the features were initially filtered by the Gain Ratio (GainRatio) approach, which is able to evaluate how each feature contributed to decreasing the overall entropy in the dataset. Secondly, principal component analysis (PCA) was used to reduce the dimensionality of the dataset into the smaller one, while still contains most of the information.

### Blastp and SCMPSP models constructions

To build a protein sequence dataset, photosynthetic proteins (6,430 sequences) and non-photosynthetic proteins (6,430 sequences) were retrieved from UniProtKB. The protein sequences were clustered to reduce sequence redundancy to lower than 25% identity before training the models. In the case of Blastp, the test dataset was used as a query to blast against the database that was built by the training dataset. A prediction class was assigned by the majority vote among the matched sequences. The best result was used to compare after varying E-value parameter. SCMPSP model was set up following the original article and the best model after the optimization process was used to compare.

## Results

### The functional relation between photosynthetic genes and their neighbors

The functional relationship between photosynthetic genes and their neighbors was analyzed in order to determine the possibility of assigning a photosynthetic function based on the neighboring gene profile. Curated gene ontology (GO) profile of photosynthetic genes of seven reference genomes (Table S2) retrieved from Uniprot was used as a curated dataset. We then investigated the relationship between the Phylo scores–a conservation score of neighboring genes–and a higher prediction specificity achieved when a more stringent criterion is used, and their functional similarity, in terms of GO. The F1 measure, which is a harmonic average between precision and recall, is used as a function-related evaluation method, and it can evaluate how much overlap there is between the GO labels of the photosynthetic queries and their neighbors. High F1 measure indicates high similarity between the GO term of a query and that of its neighbors, while low F1 measure indicates low similarity. To avoid incomplete annotation, all ancestor nodes of labeled GO terms were retrieved via GO.db, and redundant GO terms were removed.

The results of the GO profile analysis of the seven genomes were similar, with respect to trend (Figure S2). Figure 2 shows the results derived from the *Prochlorococcus Marinus* genome, as an example. We generated a random GO dataset from the pool of all GO terms annotated from all proteins in the reference genomes. The random dataset was sampled with a number equal to the number of GO terms in each conserved neighbor criteria. Without a Phylo score cutoff, the similarity between the GO terms of photosynthetic queries and their neighbors was low and almost equal to the random GO terms, probably due to the inclusion of neighboring genes that are not related to the photosynthetic function in the calculation. We found that when the Phylo score cutoff was increased, the F1 measure accordingly increased and higher than that of the random GO dataset. This result indicates that the functional relationship is not due to chance. This evidence suggests that gene neighbor conservation is a feature that can be used for functional assignment. However, we found that prediction coverage is a limitation of this method. As shown in the graph, the prediction coverage dropped continuously when the cutoff became more stringent. Therefore, it became clear that this method needed more systematic ways to guide functional assignments.

**Figure 2.**
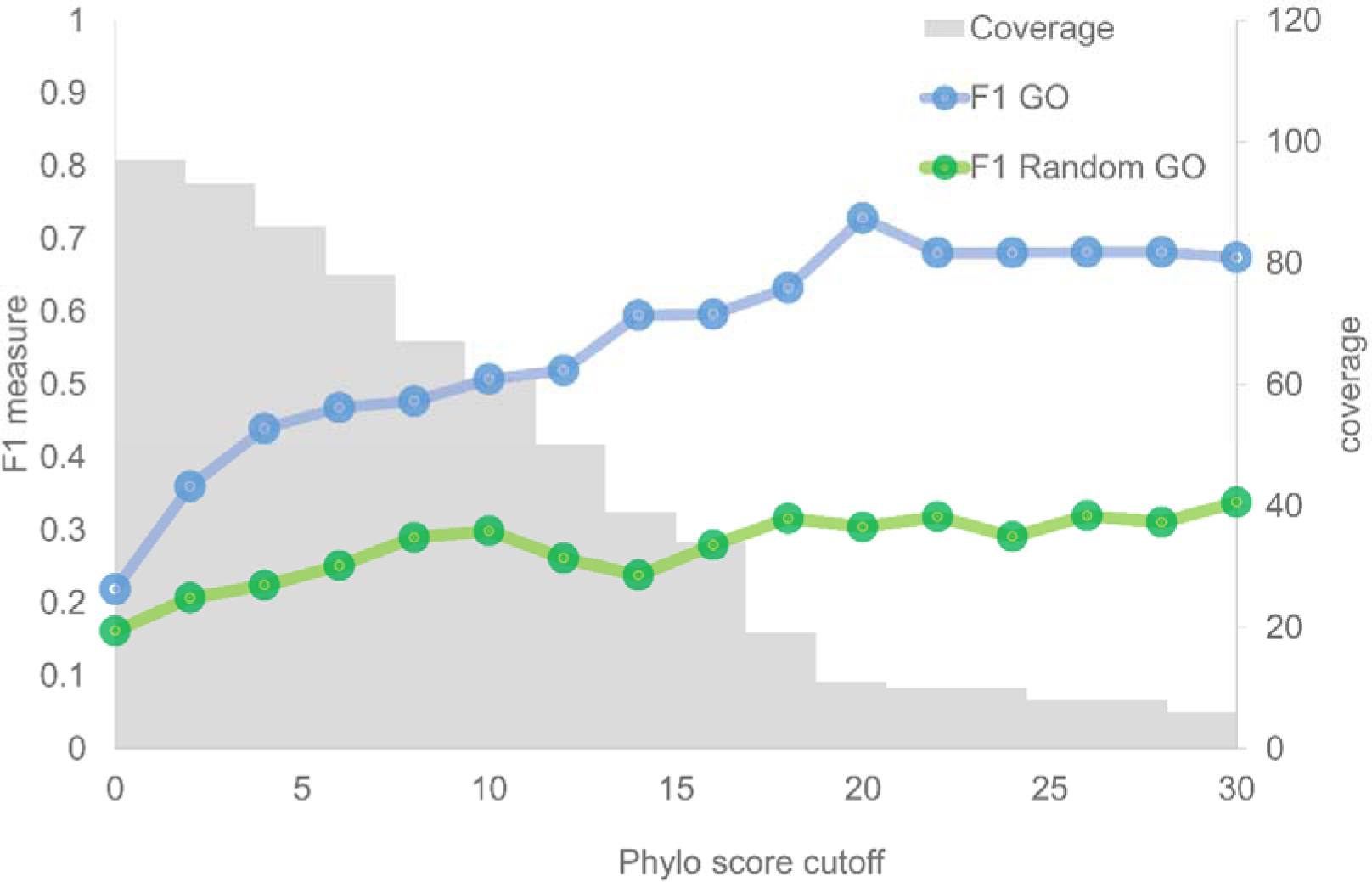
The relationship between conservation score (Phylo score) (x-axis) and F1 measure (y-axis) of *Prochlorococcus Marinus*. The blue line represents the F1 measure for indicating the similarity between GO terms from photosynthetic genes and from their neighbors. The green line represents the F1 measure for indicating the similarity between GO terms from photosynthetic genes and from random GO terms. Coverage (right x-axis) represents the number of predicted proteins.

### GNN-based machine learning model development for photosynthetic protein classification

As the Phylo scores represent the conservation of gene neighborhoods, which has the potential to refer to functional similarity, we proposed that by using the conservation profile of gene neighborhoods, photosynthetic genes/proteins can be distinguished from non-photosynthetic genes/proteins. Considering the numerous benefits of employing ML to predict protein function, the algorithms were performed using Weka software version 3.9.1 (Frank, et al., 2010). The protocol for building the model of photosynthetic gene classification namely PhotoMod (PHOTOsynthetic MODel), as shown in Figure 1, contains information on dataset building, feature extraction, data preprocessing, classifier selection and feature selection.

The performance of three different classifiers, BN, SMO and RF was addressed and compared after fine-tuning parameters (Table S3). It was found that RF showed the highest performance of up to 88% accuracy (Table 1). Also, the F1 measure of minor class and MCC indicated that RF did not suffer from the imbalanced dataset as much as the other methods, especially, BN. SMO performed well and better than BN, which was the worst classifier in all aspects. However, the average overall performance of the three classifiers, >80% of accuracy, indicates that the gene neighborhood profile can be used to discriminate between photosynthetic function and non-photosynthetic function.

**Table 1.**
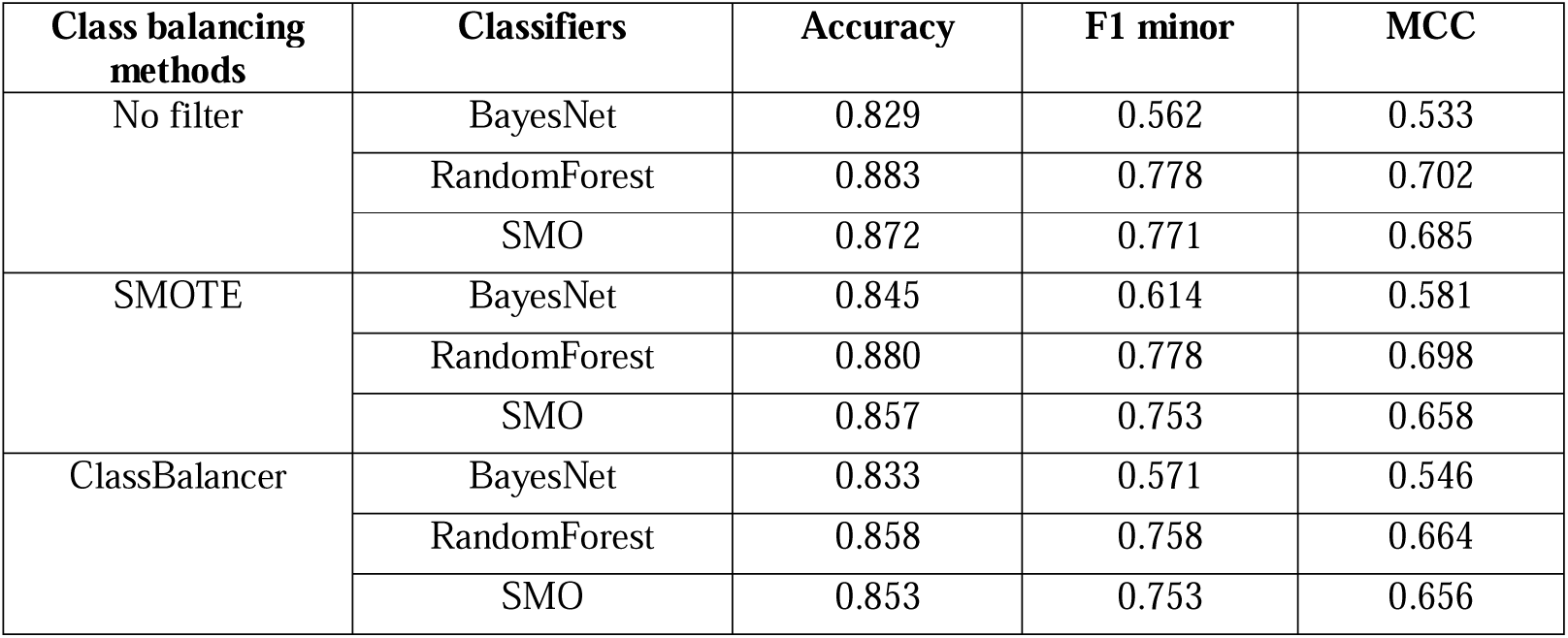
Classification performance of different classifier algorithms and class balancing methods.

To address the class imbalance problem that was found during the data preprocessing step, two supervised approaches were introduced. First, the ClassBalancer method was applied to reweight the instances of each class in the dataset equally, while the total sum of weights of all instances was maintained. The second method involved the resampling of the instances of the minor class in a dataset by applying the Synthetic Minority Oversampling TEchnique (SMOTE) (Lertampaiporn, et al., 2013). It was found that the two approaches were able to improve the prediction performance of BN, but not in SMO and RF (Table 1). The prediction performance of BN was improved in all aspects when ClassBalancer and SMOTE were applied. Both filter methods decreased the classification performance of RF classifier, which comes as no surprise because tree-based algorithms often perform well on imbalanced datasets (Galar, et al., 2012; Krawczyk, 2016). Also, it has been previously shown that many classifiers, including RF, do not benefit from SMOTE when a high-dimensional dataset is used (Blagus and Lusa, 2013), but it is not known why supervised balancing approaches struggle with high-dimensional class-imbalanced data (Blagus and Lusa, 2013). Our result suggests that RF is the most suitable approach to deal with high-dimensional class-imbalanced data without any boosting method.

In order to make the model more robust and reduce the chance of overfitting, GainRatio and PCA were applied to select optimal feature subset and reduce feature dimension, respectively, in the RF model. The result shows that the prediction performance of RF was slightly improved when the feature selection methods were applied (Figure S3). The improvement was observed when the number of selected features was at least 2,000. Decreasing the number of features below 100 features lowers performance, which might be the result of the loss of important information. The features that were selected for the model tree were considered important for photosynthesis. Although three E-value criteria were applied for protein clustering features, the most informative criterion was 1E-10 as indicated by the number of selected features (58%) as shown in Table 2. However, to achieve the highest performance, the three E-value criteria were combined. The importance of such combination was emphasized by rebuilding the model using features from single E-value criterion and their different combinations and comparing the F1 measures of the models (Table S4). The maximum performance was observed when all three E-value criteria were combined, which confirms our hypothesis that the three E-value criteria support each other, thereby enabling a better photosynthesis function classification. Although the E-value criterion of 1E-10 mainly contributed to the prediction performance, the lower E-values (1E-50 and 1E-100) are likely necessary for deeper function classification. After PCA was applied with 95% of the variance, the total number of features dropped to 301 features. The model performance was observed and compared to other methods as shown in the next section.

**Table 2.** The number of selected features for each E-value criterion.

| **E-value criteria** | **Number of selected features** |
| --- | --- |
| 1E-10 | 1,156 (58%) |
| 1E-50 | 643 (32%) |
| 1E-100 | 201 (10%) |
| Total | 2,000 |

### Performance comparison of photosynthetic function classification methods

The prediction performance of our genome neighborhood-based model (PhotoMod) was compared with those of the basic sequence similarity approach, Blastp, and advanced machine learning approach, SCMPSP. The performance of three methods was shown by using 5-folds cross-validation replicated 10 times and independent test set (Table 3). The accuracy score indicates overall correct prediction, while F1 minor is used to describe the correctness of the minor class (photosynthetic class). MCC is used to evaluate the overall model correctness in regard to its robustness to an imbalanced dataset. Blast methods achieved accuracy, F1 minor and MCC up to 0.843, 0.556 and 0.497, while SCMPSP achieved up to 0.825, 0.625 and 0.512, respectively. Notice that there was no significant difference observed between both sequence-based methods. PhotoMod overcame both methods and achieved accuracy, F1 minor and MCC up to 0.941, 0.892 and 0.852, respectively. Also, a high score of F1 minor and MCC indicates that our model does not suffer from the imbalanced dataset.

**Table 3.**
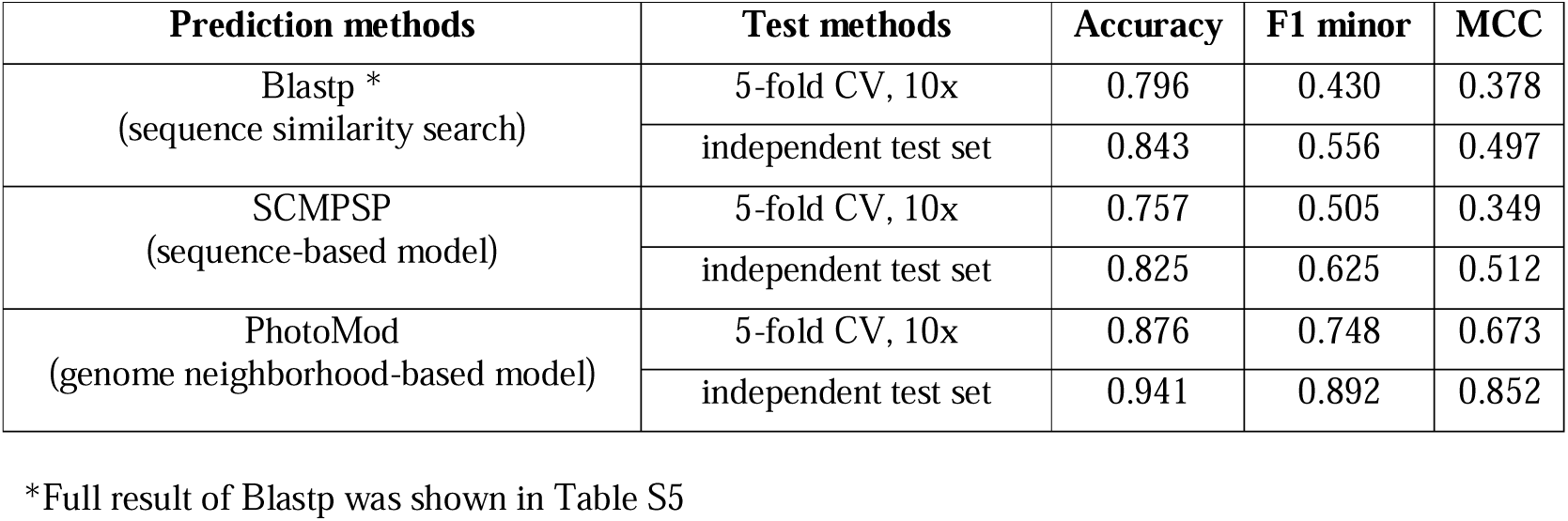
Performance comparison of different methods in the classification of photosynthetic proteins using ten times 5-fold cross-validation

### Performance comparison of photosynthetic function classification methods using novel photosynthetic protein dataset

The goodness of the PhotoMod was shown by predicting recently identified novel photosynthetic proteins and by comparison our model with other protein function prediction tools. Twelve novel photosynthetic proteins were collected from the literature. These photosynthetic proteins were recently reported and had never been deposited in any database (Table S6). To measure false positive error, we also included non-photosynthetic proteins. The non-photosynthetic dataset was collected from the recently reported prokaryotic protein sequences that contain no photosynthetic GO term label (as described in the method section), which were obtained from UniprotKB (in 2018). The protein sequences were clustered to reduce sequence redundancy to lower than 25%. In total, only 111 non-photosynthetic protein sequences passed this criterion.

The performance of the present model was compared to the other functional prediction tools that rely on amino sequence property. To enable a fair comparison, protein sequences were used as input for all prediction tools. In our case, the sequences were blasted against the protein database retrieved from the 154 proteomes. The matched sequences were used to call genome neighborhood profiles and make a prediction. A prediction class was assigned by the majority vote. The prediction performance of our model was compared with that of other prediction tools including Blastp, SCMPSP, DeepGO, and SVMprot under optimal conditions (E-value or confidence score) (Table S7).

Only three of 12 novel photosynthetic proteins could be assigned photosynthetic function by Blastp (Table 4). F1 minor (0.188) and MCC (0.078) showed that the overall performance of this method was inefficient in the prediction of photosynthesis class. However, the false positive of this method was very low, resulting in a high accuracy score (0.789). On the other hand, SVMprot predicted six proteins correctly as photosynthetic proteins and allowed higher false-positive error compared to that of Blastp. Thus, the precision of this method (0.140) was lower than the Blastp approach, but the overall performance was better (MCC = 0.104). We found that DeepGO could not be compared, because no GO term related to photosynthesis was observed. SCMPSP performed worst compared to the other methods (MCC = −0.042). Only three of 12 photosynthetic proteins were correctly predicted, while the F1 minor was only 0.120. PhotoMod showed a better predictive performance in terms of both F1 minor (0.300) and MCC score (0.214). Interestingly, the model predicted 6 of the 12 photosynthetic proteins correctly with tolerable false positive error in comparison to SVMprot and Blastp (Table S8).

**Table 4.** Performance comparison of different methods in the classification of novel photosynthetic proteins.

| **Methods** | **TP** | **TN** | **Precision** | **Recall** | **Accuracy** | **F1 minor** | **MCC** |
| --- | --- | --- | --- | --- | --- | --- | --- |
| Blastp | 3 | 94 | 0.150 | 0.250 | 0.789 | 0.188 | 0.078 |
| SVMProt | 6 | 74 | 0.140 | 0.500 | 0.650 | 0.218 | 0.104 |
| DeepGO | 0 | 111 | - | - | - | - | - |
| SCMPSP | 3 | 76 | 0.079 | 0.250 | 0.642 | 0.120 | -0.042 |
| PhotoMod | 6 | 89 | 0.214 | 0.500 | 0.772 | 0.300 | 0.214 |

The incorrect prediction results from PhotoMod were investigated further and found to be in large diverse GNNs (Figure 3). The high node variety in the three GNNs indicates that the query proteins may contribute to several pathways. The reason might be that those proteins contain 1) multiple domains, which can either contribute to one independent function or interact with other domains to do other functions (Vogel, et al., 2004), or 2) a broad function domain such as kinase domain. It has been found that RfpA has a histidine kinase domain, which acts as sensor kinase in the signal transduction cascade (Zhao, et al., 2015). DpxA contains one ubiquitous GAF domain and one histidine kinase domain for light color sensing (Bhaya, 2016). IflA consists of three GAF domains that act as photoreceptor (Bussell and Kehoe, 2013). These domains are commonly found in the non-photosynthetic function, as predicted by sequence-similarity based methods, and likely contributed to the high variation in the GNN.

**Figure 3.**
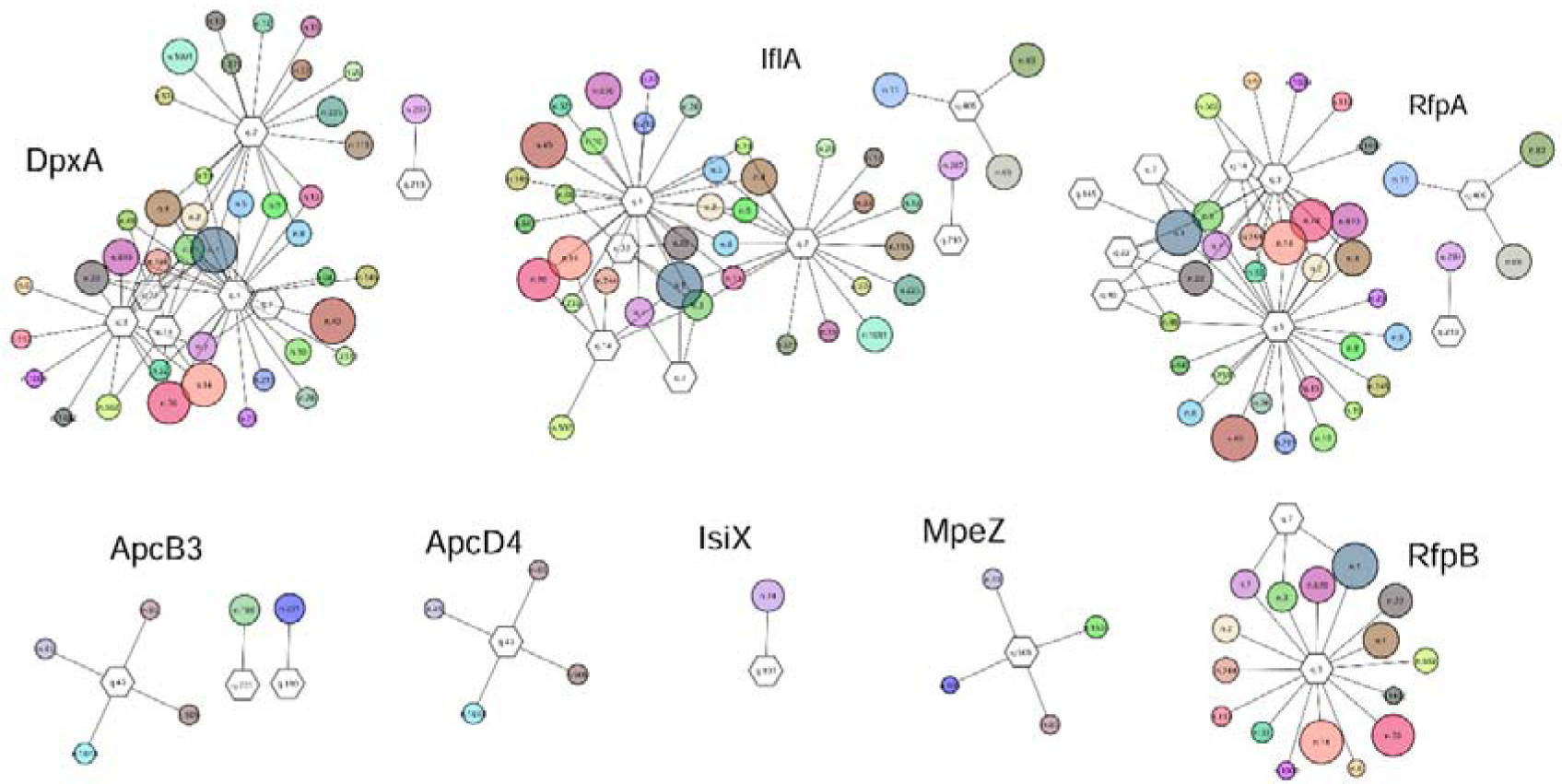
Genome neighborhood networks of novel photosynthetic genes used in this study. The protein clusters that are matched by the query sequence are represented by a hexagon. The circles represent protein cluster from the neighboring genes of which edges show gene neighborhood relationship with the query cluster. Size of the circle indicates the conservation (corresponding to Phylo score). The conserved neighboring genes are selected if the Phylo score is greater than 10. The proteins are clustered with E-value cutoff 1E-10.

### Prediction of unknown proteins in cyanobacteria genome

More than 50% of protein-coding genes in the genome of the model photosynthetic organism, *Synechocystis* sp. PCC 6803, is unknown. We applied our platform to seek for photosynthetic related genes in this organism, using the unknown protein sequences as input. Of the 1,885 unknown protein-coding genes, 479 sequences (∼26%) were found to be involved in photosynthesis (Table S9) and were validated using the information obtained from the literature. For example, the gene operon, *slr0149, slr0148, slr0147* and *slr0146* that were predicted to be related to photosynthesis, have been shown to be involved in the assembly of Photosystem II in cyanobacterium *Synechocystis* 6803 (Wegener, et al., 2008). Interestingly, it was observed that these gene set are conserved, and their expression patterns are correlated in various photosynthetic prokaryotes [unpublished data].

## Discussion

We showed that using phylogenetic distance criteria to assign protein function from gene neighborhood yields a high-sensitivity result, though the precision was very low. This particular event was also observed in the relaxed condition of gene cluster detection for functional classification (Ling, et al., 2009). It has been also shown that ML efficiently assigns protein function (Yu, et al., 2018) and predicts the gene cluster that holds functional coupling (Chuang, et al., 2012). In this study, we applied the ML approach with gene neighborhood-based feature to facilitate the photosynthetic function assignment. The ML algorithms worked well with our feature extraction scheme, and the high classification performance of all classifiers indicates the feature merit.

High-dimensional data was observed after extracting genome neighborhood information. The RF algorithm showed the highest performance, although SVM and BN have been more popular for high-dimensional data such as microarray classification (Shi, et al., 2010) and text classification (Medhat, et al., 2014). Interaction between attributes and the combination of tree predictors might be the key advantages of RF over other algorithms. It has been shown that RF performs well in the studies of interactions between genetic loci (Cordell, 2009; Moore, et al., 2010). Although RF classifier handles high-dimensional data well, redundant and useless attributes can cause time wastage for both training and testing processes. In this study, GainRatio, a feature selection method, was employed, although it ignores feature dependencies and interaction with the classifier. Nevertheless, it benefits from improved computational efficiency, enabling high-dimensional dataset analysis (Saeys, et al., 2007).

We demonstrated that our model outperforms the sequence-based methods. Basically, the sequence-based model benefits from amino acid properties that are conserved among the group of protein. However, the photosynthesis is a complex system consisting of many different proteins, therefore broad sequence property can cause low precision. Only three of 12 novel proteins could be correctly assigned photosynthetic function by Blastp, which comes as no surprise considering the difficulty of assigning photosynthetic function by sequence-similarity search (Ashkenazi, et al., 2012). We found that DeepGO could not predict any GO term that is related to photosynthesis, suggesting that DeepGO is not efficient for specific functions, e.g. photosynthesis, because it has been trained by a large number of GO terms. Also, it is reportedly not suited for predicting specific protein functions in a biological process (Kulmanov, et al., 2018). SCMPSP was shown previously that its performance was better than Blast method and comparable to the ML approach (Vasylenko, et al., 2015). However, in this study, we found that SCMPSP performed inefficiently and even worse than the Blast method. We suspect that the performance of this method depends on the genetic algorithm which might get stuck in local minima if the number of generations is not sufficiently long enough. While sequence-based models yield a low precision score, our PhotoMod combines sequence property and GNN as learning features to achieve a higher precision result. This result might indicate that photosynthetic function is governed by genomic constraints in addition to amino acid property. Nevertheless, the source of false-negative of the present method is from the diverse patterns of the GNN profile. This problem was also observed in the classification of heterogeneous data on the complex disease (Li, et al., 2018; Urbanowicz, et al., 2013). One way to deal with this problem is to reduce the number of patterns using the clustering algorithm before ML step (Li, et al., 2018).

Limitations of our approach should be also mentioned. First, it works well with only prokaryote proteins, as the proteins were extracted from only photosynthetic prokaryotes. Photosynthetic eukaryotes such as plants were excluded as the concept of gene neighborhood in eukaryotes is complicated due to their chromosome conformation (De, et al., 2009; Meadows, et al., 2010). Another limitation is that if the query gene is unique, it is impossible to find conserved neighboring genes. If the queries do not match any sequence in our database, which contains all proteins in photosynthetic organisms, we assumed that those sequences are not from a photosynthetic organism and discarded those sequences to avoid misclassification. Lastly, we found that neighborhood gene calling was a major time-consuming step of the platform. Therefore, a database scheme for collecting gene neighborhood profile is recommended to improve the performance.

## Additional Information

Supplementary data are available at http://bicep.kmutt.ac.th/photomod_standalone

## Acknowledgment

The authors thank Dr. Weerayuth Kittichotirat and Dr. Sawannee Sutheeworapong for insightful discussions and suggestions. We also thank Bioinformatics and Systems Biology Program and KMUTT to support research materials and equipment. Furthermore, we thank Mr. Oscar Nnaemeka from the School of Bioresources and Technology, KMUTT, for his editing and proofreading of the manuscript. This work was partly supported by Petchra Pra Jom Klao Doctoral Scholarship (No: 13/2558) from KMUTT and a research grant (NRMJ: 2559A30602134#60000108) from the National Research Council of Thailand (http://en.nrct.go.th). The funders had no role in study design, data collection and analysis, the decision to publish, and preparation of the manuscript.

## Author Contributions

A.S., T.L., and M.R. designed the research in the manuscript. A.S. conducted the experiments and analyzed the results. A.S., T.L., and M.R. wrote the manuscript and prepared all of the figures. All of the authors reviewed the manuscript.

## Conflict of Interest

The authors declare that they have no competing interests.

