## Supplementary material for "Photosynthetic protein classification using genome neighborhood-based machine learning feature": Text S

**SUPPLEMENTARY DATA**

**Topic:** Genome neighborhood-based machine learning feature for the classification of photosynthetic proteins

Text S 1 Collection of photosynthetic prokaryote genomes and protein classification protocol

The complete genomes of 163 photosynthetic prokaryotes, scattered across seven phyla and mostly comprising cyanobacteria, were retrieved from NCBI database. To confirm their photosynthetic ability, reaction center proteins of the photosynthetic system (Type I RC: PsaA, PsaB, PshA, PscA, Type II RC: PsbA, PsbB, PufL, PufM) were investigated. Eight of the genomes that lacked the reaction center genes and the genome of *Roseobacter litoralis*, Strain Och 149, whose reaction center genes (*pufL, pufM*) are located in the plasmid ^1^ were excluded from our analysis. Details about the remaining 154 genomes used in our study are provided in Table S 1. Although genes within the collected genomes have been identified and annotated, different gene finder programs and parameter settings were used, which might introduce differences in the number of identified genes and the quality of annotation. Therefore, we used Prodigal software as part of CMG-biotools workbench ^2^ to re-identify DNA coding regions and corresponding protein sequences from the genomes. In order to reduce sequence redundancy, Markov clustering method (MCL) ^3^ was employed to classify protein sequences into protein families ^4^.

*Protein classification protocol* MCL is an efficient tool for classifying protein families, especially with huge datasets and only requires information on sequence similarity relationships, which is generally obtained using BLASTP with the all-vs-all protocol. The BLASTP all-vs-all comparison was performed on the combined all proteins in a FASTA file that was obtained from a public database or was newly called from genomes by prediction tools. BLASTP standalone version was carried out with the NCBI BLAST package v2.2.31. All-vs-all comparison is achieved by using the combined FASTA file searching against database of themselves. Making database of the combined FASTA file is as follows:

*makeblastdb -in all_proteins.fasta -parse_seqids -dbtype prot*

BLASTP was applied with default parameters and output format number 6. The maximum number of HSPs per subject sequence was set to 1. E-value cutoff can be varied to different values (in this study 1E-10, 1E-50 and 1E-100). The command is shown below where [X] is adjusted e-value:

*blastp -db all_proteins.fasta -query all_proteins.fasta  -out all_proteins.blastout -max_hsps 1 -outfmt '6' -evalue [X] -num_threads 8*

MCL version 14-137 was obtained from http://www.micans.org/mcl/. MCL accepts sequence similarity information in ‘ABC’ format, which is a three-column file containing qseqid, sseqid and e-value. We can simply generate this file via standard Unix command as following.

*cut -f 1,2,11 all_proteins.blastout > all_proteins.abc*

The abc-format file “all_proteins.abc” is then executed by mcxload, which creates two output files, a network file called “all_proteins.mci” and a label information file in “all_proteins.dict”. The option --stream-mirror option is applied to enforce undirected graph. The option --stream-neg-log10 transform e-value input to log-10 representation. The maximum e-value is set by last option -stream-tf 'ceil(200)', which any E-value below 1e-200 is set to a maximum allowed edge weight of 200.

*mcxload -abc all_proteins.abc -write-tab all_proteins.dict -o all_proteins.mci --stream-mirror --stream-neg-log10 -stream-tf 'ceil(200)'*

Now, MCL is ready to run by using all_proteins.mci as input. Only one parameter can be adjusted is inflation values. The higher inflation value yields the higher clustering quality. Output file is automatically created to out.all_proteins.mci.I[Y], where [Y] is inflation value.

*mcl all_proteins.mci -I [Y]*

Then, the raw output can be converted to a readable format with labels by mcxdump. The output file contains all proteins for a cluster on a single line delimited by tab.

*mcxdump -icl out.all_proteins.mci.I[Y] -o dump.all_proteins.mci.I[Y] –tabr* all_proteins.dict

Text S 2 Identification of photosynthetic genes using photosynthesis-specific GO terms

### The protein sequences that contain at least one of 61 photosynthesis-specific GO terms reported by Ashkenazi et al. ^5^ were collected as an initial positive dataset. In total, 15,195 unique proteins labeled with at least one of 61 GO terms were found, confirming their photosynthetic function. The photosynthetic function of initial positive dataset was transferred to our protein dataset, which is a complete set of proteomes of 154 photosynthetic prokaryote genomes. The photosynthetic function was transferred if a percent identity and percent sequence coverage of more than 80% were obtained. Finally, the positive data set contained 6,430 protein sequences. The negative dataset was collected from non-photosynthetic genes in UniprotKB. The protein sequences from UniprotKB that were not labeled with any photosynthetic GO term were randomly selected. To ensure the photosynthetic function is not present in the negative data set, all ancestor nodes of annotated GO terms in each sequence were retrieved using GO.db library. Sequences containing GO nodes that match the photosynthetic function were removed. Additionally, the sequences are necessarily annotated at least to GO level 3, which is the same level of photosynthesis (GO:0015979), otherwise they were automatically removed. The sequences that passed the criteria were blasted against our protein dataset with stringent criteria (percent identity and coverage >80%) ^6^. The final non-photosynthetic dataset was randomly selected to be equal to the number of positive dataset from the pool of matched protein sequences.

Text S 3 Evaluation metrics

We selected and used three different metrics to evaluate the prediction performance of classifiers. Accuracy ^7^ is the most common metric for classifier evaluation. It measures the overall performance by considering the probability of true value among the predictions.

Accuracy = $\frac{TP+TN}{TP+TN+FP+FN}$

F1 measure ^7^ is the harmonic mean of precision and recall. Precision is a measurement of correctness i.e. how many examples of the positive class were correctly predicted among the total positive predictions. Recall is a measurement of completeness i.e. how many examples of the positive class were correctly predicted among the total examples in the positive class. We measure the F1 score of the minor class (photosynthetic class) to measure the real performance of the class-imbalanced dataset.

F1 measure = $\frac{2 x Precision x Recall}{Precision x Recall}$

Precision = $\frac{TP}{TP+ FP}$

Recall = $\frac{TP}{TP+ FN}$

Matthews Correlation Coefficient ^8,9^ is a single formula measurement metric that considers mutually accuracies and error rates on both classes. It is widely used in bioinformatics area because of its robustness in an imbalanced dataset.

MCC = $\frac{(TP x TN)-(FP x FN)}{\sqrt{(TP+FP)(TP+FN)(TN+FP)(TN+FN)}}$

The best achievable score of each classifier after varying threshold is used to be a final score for model comparison.

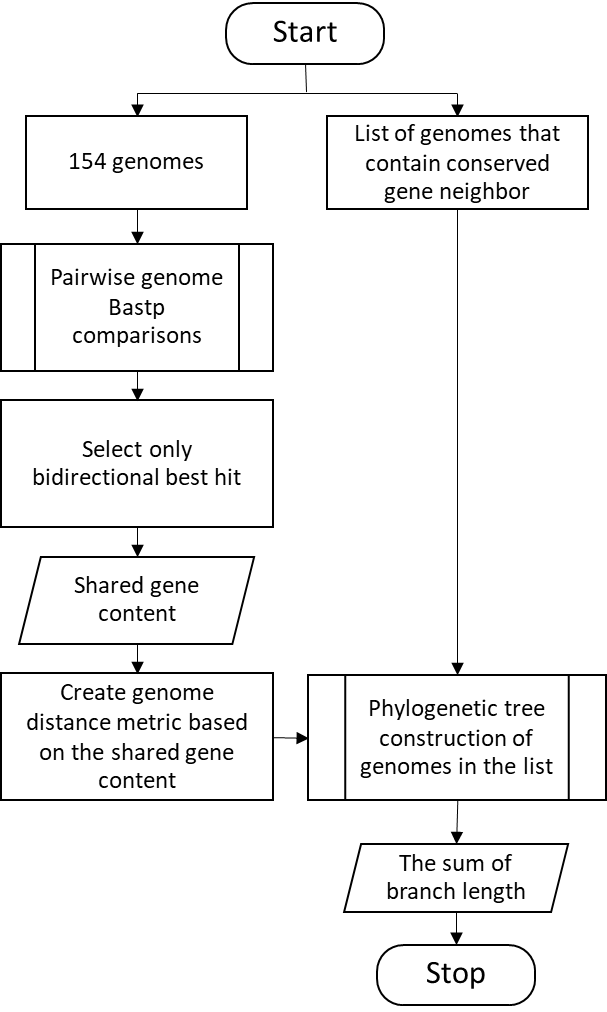

**Figure S 1** Flowchart of Phylo score calculation

| *Chlorobaculum tepidum* | *Chloroflexus aurantiacus* |
| --- | --- |
| 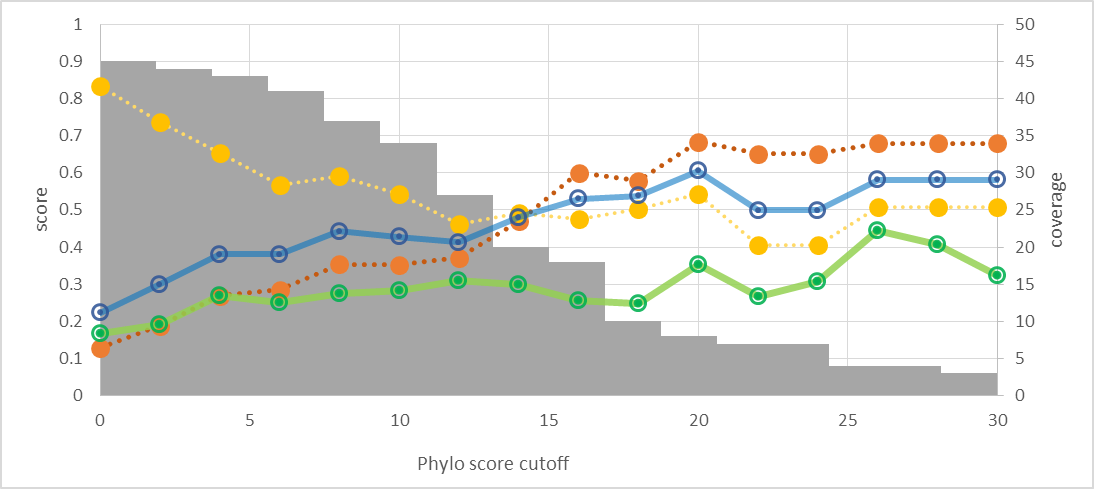 | 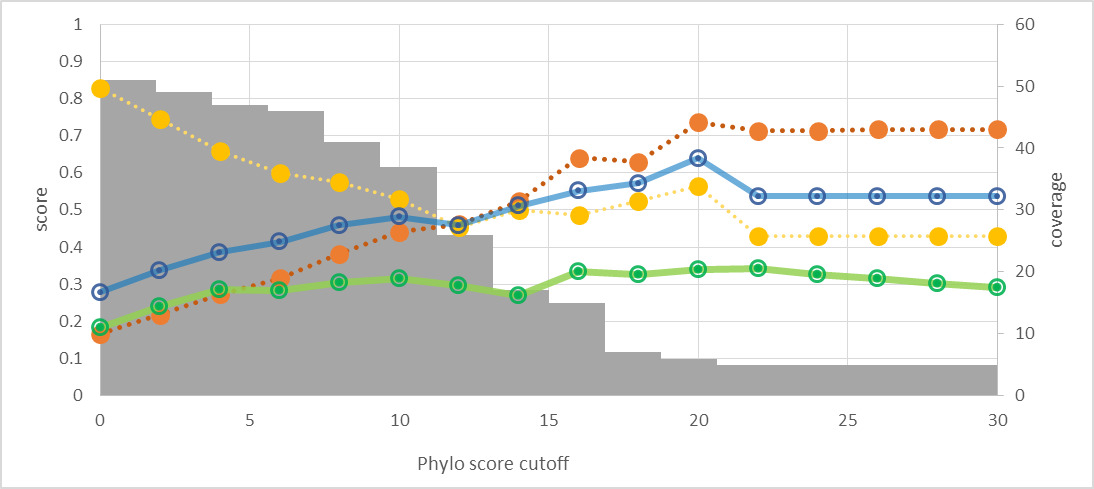 |
| *Rhodobacter sphaeroides* | *Rhodospirillum rubrum* |
| 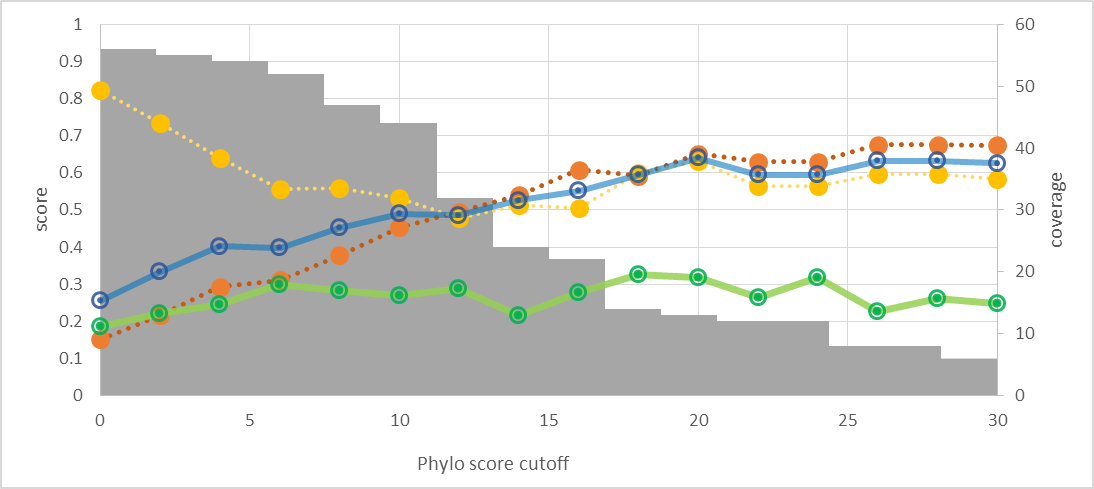 | 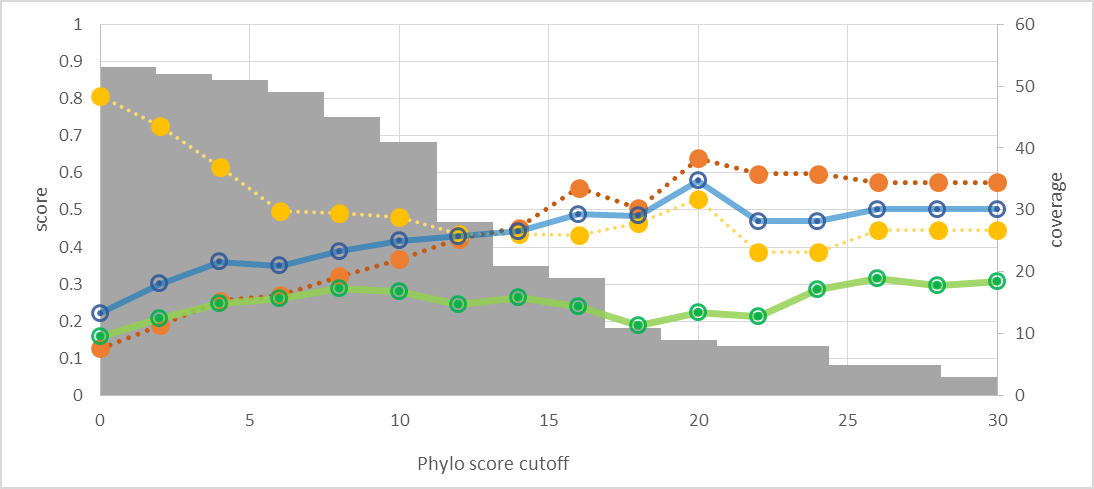 |
| *Prochlorococcus marinus* | *Thermosynechococcus elongatus* |
| 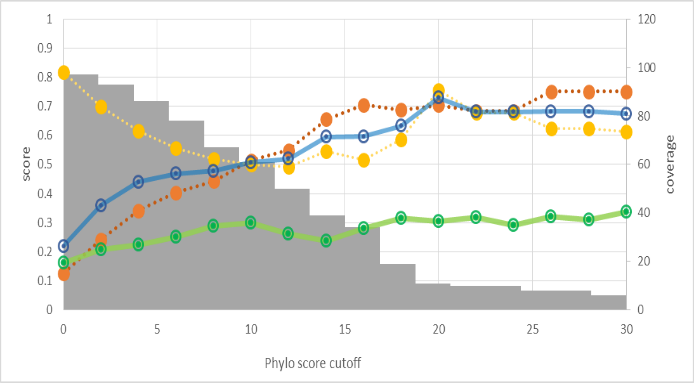 | 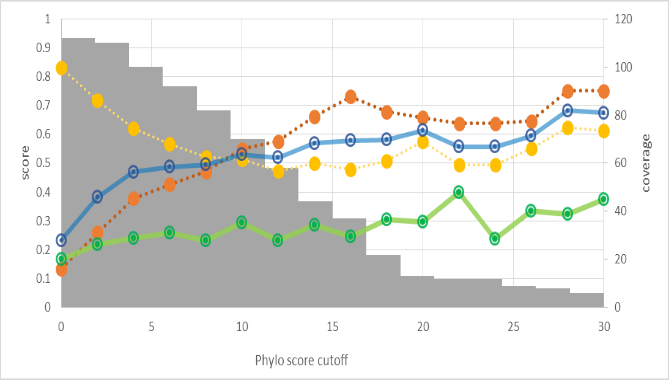 |
| *Gloeobacter violaceus* | |
| 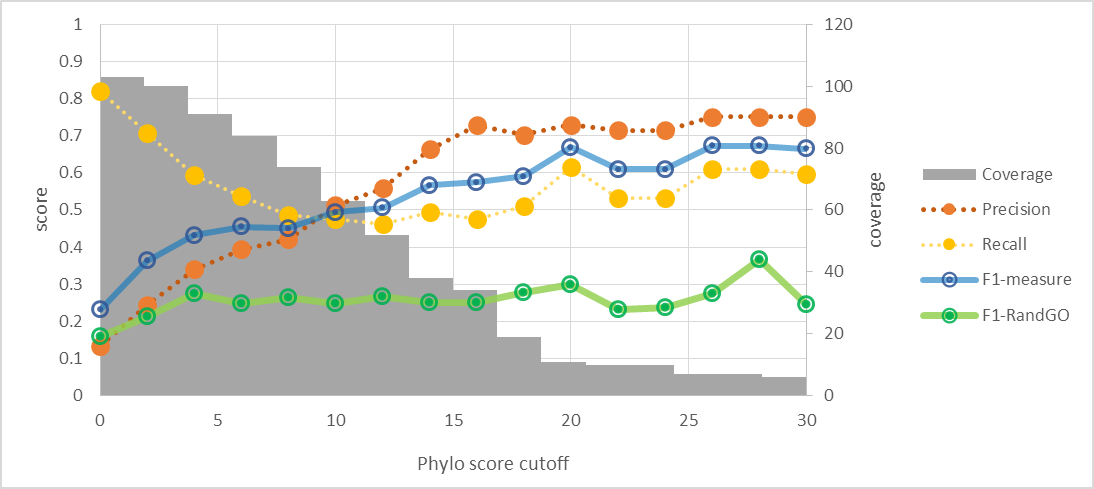 | |
| **Figure S 2** Functional relation measurements between photosynthetic genes and their neighbors using Phylo score as a cutoff criteria | |

Evaluation score

| Number of features | Accuracy | F1 minor | MCC |
| --- | --- | --- | --- |
| 100 | 0.8494 | 0.645 | 0.593 |
| 500 | 0.88955 | 0.762 | 0.709 |
| 1000 | 0.8949 | 0.779 | 0.724 |
| 2000 | 0.89924 | 0.81 | 0.744 |
| 3000 | 0.89822 | 0.81 | 0.743 |
| 4000 | 0.89787 | 0.812 | 0.743 |
| 5000 | 0.89677 | 0.811 | 0.741 |
| 6000 | 0.89821 | 0.808 | 0.741 |
| 7000 | 0.90111 | 0.811 | 0.747 |
| 8000 | 0.89618 | 0.806 | 0.738 |
| 9000 | 0.89642 | 0.806 | 0.738 |
| 10000 | 0.89626 | 0.804 | 0.736 |
| 11000 | 0.89558 | 0.803 | 0.735 |
| 12000 | 0.89583 | 0.804 | 0.736 |
| 13000 | 0.89609 | 0.805 | 0.737 |
| W/O FS | 0.89464 | 0.804 | 0.735 |

**Figure S 3** Minimization of the number of the feature using GainRatio method and random forest classifier

**Table S 1** List of 154 complete genomes of photosynthetic prokaryotes after filtering by reaction center detection

| **ID** | **Taxonomy** | **Organism** | **NCBI Accesion** |
| --- | --- | --- | --- |
| 1 | Acidobacteria | Candidatus Chloracidobacteriumthermophilum_1 | CP002514 |
| 1 | Acidobacteria | Candidatus Chloracidobacteriumthermophilum_2 | CP002515 |
| 5 | Chlorobi (Green sulfur bacteria) | Chlorobium tepidum | AE006470 |
| 6 | Chlorobi (Green sulfur bacteria) | Chlorobaculum parvum | CP001099 |
| 7 | Chlorobi (Green sulfur bacteria) | Chlorobium limicola | CP001097 |
| 8 | Chlorobi (Green sulfur bacteria) | Chloroherpeton thalassium ATCC 35110 | CP001100 |
| 9 | Chlorobi (Green sulfur bacteria) | Prosthecochloris aestuarii | CP001108 |
| 10 | Chlorobi (Green sulfur bacteria) | Chlorobium phaeovibrioides DSM 265 | CP000607 |
| 11 | Chlorobi (Green sulfur bacteria) | Chlorobium phaeobacteroides BS1 | CP001101 |
| 12 | Chlorobi (Green sulfur bacteria) | Chlorobium chlorochromatii CaD3 | CP000108 |
| 13 | Chlorobi (Green sulfur bacteria) | Pelodictyon phaeoclathratiforme BU-1 | CP001110 |
| 14 | Chlorobi (Green sulfur bacteria) | Chlorobium luteolum DSM 273 | CP000096 |
| 15 | Chlorobi (Green sulfur bacteria) | Prosthecochloris sp. CIB 2401 | CP016432.1 |
| 16 | Chlorobi (Green sulfur bacteria) | Chlorobium phaeobacteroides DSM 266 | CP000492.1 |
| 17 | Chloroflexi (Green nonsulfur bacteria) | Chloroflexus aurantiacus | CP000909 |
| 18 | Chloroflexi (Green nonsulfur bacteria) | Roseiflexus castenholzii | CP000804 |
| 20 | Chloroflexi (Green nonsulfur bacteria) | Roseiflexus sp. RS-1 | CP000686 |
| 21 | Chloroflexi (Green nonsulfur bacteria) | Chloroflexus aggregans DSM 9485 | CP001337 |
| 22 | Chloroflexi (Green nonsulfur bacteria) | Chloroflexus sp. Y-400-fl | CP001364 |
| 23 | Firmicutes (Heliobacteria) | Heliobacterium modesticaldum Ice1 | CP000930 |
| 25 | Proteobacteria (Purple bacteria) | Allochromatium vinosum | CP001896 |
| 26 | Proteobacteria (Purple bacteria) | Bradyrhizobium sp. BTAi1 | CP000494 |
| 27 | Proteobacteria (Purple bacteria) | Bradyrhizobium sp. ORS 278 | CU234118 |
| 28 | Proteobacteria (Purple bacteria) | Congregibacter litoralis KT71 | CM002299 |
| 29 | Proteobacteria (Purple bacteria) | Dinoroseobacter shibae DFL 12 = DSM 16493 | CP000830 |
| 30 | Proteobacteria (Purple bacteria) | Jannaschia sp. CCS1 | CP000264 |
| 34 | Proteobacteria (Purple bacteria) | Methylobacterium extorquens PA1 | CP000908 |
| 35 | Proteobacteria (Purple bacteria) | Methylobacterium radiotolerans JCM 2831 | CP001001 |
| 36 | Proteobacteria (Purple bacteria) | Rhodobacter capsulatus | CP001312 |
| 37 | Proteobacteria (Purple bacteria) | Rhodobacter sphaeroides 2.4.1_1 | CP000143 |
| 37 | Proteobacteria (Purple bacteria) | Rhodobacter sphaeroides 2.4.1_2 | CP000144 |
| 39 | Proteobacteria (Purple bacteria) | Rhodobacter sphaeroides ATCC 17025 | CP000661 |
| 40 | Proteobacteria (Purple bacteria) | Rhodobacter sphaeroides ATCC 17029_1 | CP000577 |
| 40 | Proteobacteria (Purple bacteria) | Rhodobacter sphaeroides ATCC 17029_2 | CP000578 |
| 42 | Proteobacteria (Purple bacteria) | Rhodopseudomonas palustris BisA53 | CP000463 |
| 43 | Proteobacteria (Purple bacteria) | Rhodopseudomonas palustris BisB18 | CP000301 |
| 44 | Proteobacteria (Purple bacteria) | Rhodopseudomonas palustris BisB5 | CP000283 |
| 45 | Proteobacteria (Purple bacteria) | Rhodopseudomonas palustris CGA009 | BX571963 |
| 46 | Proteobacteria (Purple bacteria) | Rhodopseudomonas palustris HaA2 | CP000250 |
| 47 | Proteobacteria (Purple bacteria) | Rhodospirillum rubrum | CP000230 |
| 48 | Proteobacteria (Purple bacteria) | Roseobacter denitrificans OCh 114 | CP000362 |
| 50 | Proteobacteria (Purple bacteria) | Rubrivivax gelatinosus IL144 | AP012320 |
| 51 | Proteobacteria (Purple bacteria) | Thiocystis violascens | CP003154 |
| 52 | Proteobacteria (Purple bacteria) | Halorhodospira halophila SL1 | CP000544 |
| 53 | Proteobacteria (Purple bacteria) | Citromicrobium sp. JL477 | CP011344.1 |
| 55 | Proteobacteria (Purple bacteria) | Acidiphilium multivorum AIU301 | AP012035 |
| 56 | Proteobacteria (Purple bacteria) | Bradyrhizobium sp. S23321 | AP012279 |
| 57 | Proteobacteria (Purple bacteria) | Brevundimonas subvibrioides ATCC15264 | CP002102 |
| 58 | Proteobacteria (Purple bacteria) | Methylobacterium chloromethanicum CM4 | CP001298 |
| 59 | Proteobacteria (Purple bacteria) | Methylobacterium extorquens AM1 | CP001510 |
| 60 | Proteobacteria (Purple bacteria) | Methylobacterium extorquens DM4 | FP103042 |
| 61 | Proteobacteria (Purple bacteria) | Methylobacterium populi BJ001 | CP001029 |
| 62 | Proteobacteria (Purple bacteria) | Methylobacterium sp. 4–46 | CP000943 |
| 63 | Proteobacteria (Purple bacteria) | Methylocella silvestris BL2 | CP001280 |
| 64 | Proteobacteria (Purple bacteria) | Rhodobacter sphaeroides KD131 | CP001150 |
| 65 | Proteobacteria (Purple bacteria) | Rhodocista centenaria SW | CP000613 |
| 66 | Proteobacteria (Purple bacteria) | Rhodomicrobium vannielii ATCC17100 | CP002292 |
| 67 | Proteobacteria (Purple bacteria) | Rhodopseudomonas palustris DX-1 | CP002418 |
| 68 | Proteobacteria (Purple bacteria) | Rhodopseudomonas palustris TIE-1 | CP001096 |
| 69 | Proteobacteria (Purple bacteria) | Rhodospirillum photometricum DSM 122 | HE663493 |
| 70 | Proteobacteria (Purple bacteria) | Rhodospirillum rubrum F11 | CP003046 |
| 71 | Proteobacteria (Purple bacteria) | Marichromatium purpuratum 984 | CP007031 |
| 72 | Gemmatimonadetes | Gemmatimonas phototrophica strain AP64 | CP011454 |
| 73 | Cyanobacteria | Prochlorococcus marinus subsp. marinus str. CCMP1375 | AE017126.1 |
| 74 | Cyanobacteria | Synechococcus elongatus PCC 7942 | CP000100.1 |
| 75 | Cyanobacteria | Microcystis aeruginosa NIES-843 | AP009552.1 |
| 76 | Cyanobacteria | Nostoc punctiforme PCC 73102 | CP001037.1 |
| 77 | Cyanobacteria | Thermosynechococcus elongatus BP-1 | BA000039.2 |
| 78 | Cyanobacteria | Trichodesmium erythraeum IMS101 | CP000393.1 |
| 79 | Cyanobacteria | Gloeobacter violaceus PCC 7421 | BA000045.2 |
| 80 | Cyanobacteria | Acaryochloris marina MBIC11017 | CP000828.1 |
| 81 | Cyanobacteria | Anabaena variabilis ATCC 29413 | CP000117.1 |
| 82 | Cyanobacteria | 'Nostoc azollae' 0708 | CP002059.1 |
| 83 | Cyanobacteria | Chroococcidiopsis thermalis PCC 7203 | CP003597.1 |
| 84 | Cyanobacteria | Cyanobacterium stanieri PCC 7202 | CP003940.1 |
| 85 | Cyanobacteria | Anabaena cylindrica PCC 7122 | CP003659.1 |
| 86 | Cyanobacteria | Cyanobium gracile PCC 6307 | CP003495.1 |
| 87 | Cyanobacteria | Dactylococcopsis salina PCC 8305 | CP003944.1 |
| 88 | Cyanobacteria | Oscillatoria acuminata PCC 6304 | CP003607.1 |
| 89 | Cyanobacteria | Crinalium epipsammum PCC 9333 | CP003620.1 |
| 90 | Cyanobacteria | Stanieria cyanosphaera PCC 7437 | CP003653.1 |
| 91 | Cyanobacteria | Synechococcus sp. CC9902 | CP000097.1 |
| 92 | Cyanobacteria | Nostoc sp. PCC 7120 | BA000019.2 |
| 93 | Cyanobacteria | Synechocystis sp. PCC 6803 | BA000022.2 |
| 94 | Cyanobacteria | Calothrix sp. 336/3 | CP011382.1 |
| 95 | Cyanobacteria | Pseudanabaena sp. PCC 7367 | CP003592.1 |
| 96 | Cyanobacteria | Anabaena sp. 90_1 | CP003284.1 |
| 96 | Cyanobacteria | Anabaena sp. 90_2 | CP003285.1 |
| 98 | Cyanobacteria | Geitlerinema sp. PCC 7407 | CP003591.1 |
| 99 | Cyanobacteria | Pleurocapsa sp. PCC 7327 | CP003590.1 |
| 100 | Cyanobacteria | Microcoleus sp. PCC 7113 | CP003630.1 |
| 101 | Cyanobacteria | Gloeocapsa sp. PCC 7428 | CP003646.1 |
| 102 | Cyanobacteria | Cyanothece sp. ATCC 51142_1 | CP000806.1 |
| 102 | Cyanobacteria | Cyanothece sp. ATCC 51142_2 | CP000807.1 |
| 104 | Cyanobacteria | Prochlorococcus sp. MIT 0801 | CP007754.1 |
| 105 | Cyanobacteria | Halothece sp. PCC 7418 | CP003945.1 |
| 106 | Cyanobacteria | Rivularia sp. PCC 7116 | CP003549.1 |
| 107 | Cyanobacteria | Cyanobacterium aponinum PCC 10605 | CP003947.1 |
| 108 | Cyanobacteria | Oscillatoria nigro-viridis PCC 7112 | CP003614.1 |
| 109 | Cyanobacteria | Gloeobacter kilaueensis JS1 | CP003587.1 |
| 111 | Cyanobacteria | Microcystis panniformis FACHB-1757 | CP011339.1 |
| 112 | Cyanobacteria | Geminocystis sp. NIES-3708 | AP014815.1 |
| 113 | Cyanobacteria | Geminocystis sp. NIES-3709 | AP014821.1 |
| 114 | Cyanobacteria | Cyanothece sp. PCC 8801 | CP001287.1 |
| 115 | Cyanobacteria | Fischerella sp. NIES-3754 | AP017305.1 |
| 116 | Cyanobacteria | Calothrix sp. PCC 7507 | CP003943.1 |
| 117 | Cyanobacteria | Nostoc sp. PCC 7524 | CP003552.1 |
| 118 | Cyanobacteria | Synechococcus sp. JA-3-3Ab | CP000239.1 |
| 119 | Cyanobacteria | Microcystis aeruginosa NIES-2549 | CP011304.1 |
| 120 | Cyanobacteria | Synechococcus elongatus PCC 6301 | AP008231.1 |
| 121 | Cyanobacteria | Prochlorococcus marinus subsp. pastoris str. CCMP1986 | BX548174.1 |
| 122 | Cyanobacteria | Prochlorococcus marinus str. MIT 9313 | BX548175.1 |
| 123 | Cyanobacteria | Synechococcus sp. JA-2-3B'a(2-13) | CP000240.1 |
| 124 | Cyanobacteria | Nostoc sp. PCC 7107 | CP003548.1 |
| 125 | Cyanobacteria | Synechocystis sp. PCC 6803 | AP012205.1 |
| 126 | Cyanobacteria | Calothrix sp. PCC 6303 | CP003610.1 |
| 127 | Cyanobacteria | Anabaena sp. wa102 | CP011456.1 |
| 128 | Cyanobacteria | Cyanothece sp. PCC 7424 | CP001291.1 |
| 129 | Cyanobacteria | Prochlorococcus sp. MIT 0604 | CP007753.1 |
| 130 | Cyanobacteria | Cyanothece sp. PCC 7425 | CP001344.1 |
| 131 | Cyanobacteria | Synechocystis sp. PCC 6803 substr. GT-I | AP012276.1 |
| 132 | Cyanobacteria | Nostoc sp. NIES-3756 | AP017295.1 |
| 133 | Cyanobacteria | Synechococcus sp. CC9311 | CP000435.1 |
| 134 | Cyanobacteria | Prochlorococcus marinus str. NATL2A | CP000095.2 |
| 135 | Cyanobacteria | Prochlorococcus marinus str. MIT 9312 | CP000111.1 |
| 136 | Cyanobacteria | Synechococcus sp. PCC 7002 | CP000951.1 |
| 137 | Cyanobacteria | Leptolyngbya sp. PCC 7376 | CP003946.1 |
| 138 | Cyanobacteria | Synechocystis sp. PCC 6803 substr. PCC-N | AP012277.1 |
| 139 | Cyanobacteria | Cyanothece sp. PCC 7822 | CP002198.1 |
| 140 | Cyanobacteria | Synechocystis sp. PCC 6803 substr. PCC-P | AP012278.1 |
| 141 | Cyanobacteria | Prochlorococcus marinus str. AS9601 | CP000551.1 |
| 142 | Cyanobacteria | Prochlorococcus marinus str. MIT 9515 | CP000552.1 |
| 143 | Cyanobacteria | Synechococcus sp. WH 7803 | CT971583.1 |
| 144 | Cyanobacteria | Synechocystis sp. PCC 6803 | CP003265.1 |
| 145 | Cyanobacteria | Cyanothece sp. PCC 8802 | CP001701.1 |
| 146 | Cyanobacteria | Synechocystis sp. PCC 6714 | CP007542.1 |
| 147 | Cyanobacteria | Leptolyngbya sp. O-77 | AP017367.1 |
| 148 | Cyanobacteria | Synechococcus sp. RCC307 | CT978603.1 |
| 149 | Cyanobacteria | Prochlorococcus marinus str. MIT 9301 | CP000576.1 |
| 150 | Cyanobacteria | Prochlorococcus marinus str. MIT 9215 | CP000825.1 |
| 151 | Cyanobacteria | Leptolyngbya sp. NIES-3755 | AP017308.1 |
| 152 | Cyanobacteria | Synechocystis sp. PCC 6803 | CP012832.1 |
| 153 | Cyanobacteria | Prochlorococcus marinus str. NATL1A | CP000553.1 |
| 154 | Cyanobacteria | Prochlorococcus marinus str. MIT 9303 | CP000554.1 |
| 155 | Cyanobacteria | Synechococcus sp. KORDI-100 | CP006269.1 |
| 156 | Cyanobacteria | Synechococcus sp. KORDI-49 | CP006270.1 |
| 157 | Cyanobacteria | Synechococcus sp. PCC 6312 | CP003558.1 |
| 158 | Cyanobacteria | Synechococcus sp. PCC 7502 | CP003594.1 |
| 159 | Cyanobacteria | Synechococcus sp. CC9605 | CP000110.1 |
| 160 | Cyanobacteria | Synechococcus sp. WH 8109 | CP006882.1 |
| 161 | Cyanobacteria | Synechococcus sp. UTEX 2973 | CP006471.1 |
| 162 | Cyanobacteria | Synechococcus sp. WH 8103 | LN847356.1 |
| 163 | Cyanobacteria | Synechococcus sp. PCC 73109 | CP013998.1 |
| 164 | Cyanobacteria | Synechococcus sp. WH 8102 | BX548020.1 |
| 165 | Cyanobacteria | Microcystis aeruginosa NIES-2481 | CP012375.1 |
| 166 | Cyanobacteria | Synechococcus sp. KORDI-52 | CP006271.1 |
| 167 | Cyanobacteria | Synechococcus sp. PCC 7003 | CP016474.1 |
| 168 | Cyanobacteria | Synechococcus sp. PCC 7117 | CP016477.1 |
| 169 | Cyanobacteria | Synechococcus sp. PCC 8807 | CP016483.1 |

**Table S 2** List of seven photosynthetic reference genomes

| No. | Organism name | Chromosome ID (GenBank) | Phylum | Photosynthetic protein found  ( Total = 241) |
| --- | --- | --- | --- | --- |
| 1 | *Chlorobaculum tepidum* | AE006470.1 | Chlorobi | 63 |
| 2 | *Chloroflexus aurantiacus* | CP000909.1 | Chloroflexi | 74 |
| 3 | *Gloeobacter violaceus* | BA000045.2 | Cyanobacteria | 178 |
| 4 | *Prochlorococcus marinus* | AE017126.1 | Cyanobacteria | 159 |
| 5 | *Rhodobacter sphaeroides* | CP000144.2,CP000143.2 | Proteobacteria | 77 |
| 6 | *Rhodospirillum rubrum* | CP000230.1 | Proteobacteria | 78 |
| 7 | *Thermosynechococcus elongatus* | BA000039.2 | Cyanobacteria | 193 |

**Table S 3** Parameter setting for each classifier

| classifier | Parameter setting after fine-tuning | Tuned parameter (range) |
| --- | --- | --- |
| BayesNet | Estimator algorithm = SimpleEstimator -A 0.1  searchAlgorithm = hill climbing algorithm (K2) | Alpha = 0.1 – 0.9 |
| SMO | calibration method = Logistic  kernel = RBFKernel  Gamma (-G) = 0.01  C parameter (-C) = 10 | Gamma = 0.001 - 10  C = 0.01 - 1000 |
| RandomForest | numIterations (number of trees) = 100  maxDepth = 0 (unlimited)  numFeatures (randomly chosen attributes) = 100 | Number of tree = 1 - 10000  Randomly chosen attributes = 1 - 10000 |

**Table S 4** Random forest classifier performance with different combinations of E-value criteria

| E-value criteria | Accuracy | F1 minor | MCC |
| --- | --- | --- | --- |
| E10+E50+E100 | 89.924(1.425) | 0.810(0.026) | 0.744(0.034) |
| E10+E50 | 86.754(1.817) | 0.710(0.053) | 0.630(0.058) |
| E10+E100 | 87.866(2.128) | 0.780(0.039) | 0.697(0.053) |
| E50+E100 | 88.224(1.881) | 0.709(0.050) | 0.650(0.058) |
| E10 | 79.382(2.698) | 0.570(0.067) | 0.440(0.079) |
| E50 | 83.954(2.304) | 0.467(0.081) | 0.396(0.091) |
| E100 | 84.034(2.339) | 0.572(0.078) | 0.509(0.083) |

**Table S 5** Prediction performance of Blastp model with different E-value parameter (done by 10 times of 5-fold cross-validation)

| E-value | F1 | Accuracy | MCC |
| --- | --- | --- | --- |
| 0.0001 | 0.406 | 0.578 | 0.135 |
| 0.001 | 0.424 | 0.586 | 0.162 |
| 0.01 | 0.444 | 0.604 | 0.196 |
| 0.1 | 0.461 | 0.634 | 0.231 |
| 1 | 0.546 | 0.732 | 0.374 |
| **10** | **0.430** | **0.796** | **0.378** |
| 100 | 0.007 | 0.748 | 0.022 |

Table S 6 List of 12 novel genes in photosystem

| Gene name | Function | Ref |
| --- | --- | --- |
| rfpA | control the expression of the Far-red light photoacclimation (FaRLiP) gene cluster | ^11^ |
| rfpB | control the expression of the Far-red light photoacclimation (FaRLiP) gene cluster | ^11^ |
| IflA | photoreceptor and regulator of cyanobacteriochrome (CBCR) (influenced by far-red light) | ^12^ |
| DpxA | photoreceptor and regulator of cyanobacteriochrome (CBCR) (represses phycoerythrin accumulation in yellow light (570–590 nm)) | ^13^ |
| fciA | transcriptional regulators responsible for tuning the phycourobilin:phycoerythrobilin ratio in response to BL and GL | ^14^ |
| fciB | transcriptional regulators responsible for tuning the phycourobilin:phycoerythrobilin ratio in response to BL and GL | ^14^ |
| isiX | a specialized antenna protein that function in *Synechococcus* PE, A4 and A14 strains under low irradiance or possibly far-red light (or both) conditions | ^15^ |
| apcD4 | a specialized antenna protein that function in *Synechococcus* PE, A4 and A14 strains under low irradiance or possibly far-red light (or both) conditions | ^15^ |
| apcB3 | a specialized antenna protein that function in *Synechococcus* PE, A4 and A14 strains under low irradiance or possibly far-red light (or both) conditions | ^15^ |
| MpeZ | a phycoerythrin-specific bilin lyase, contributes to the type IV chromatic acclimation (CA4) response by attaching phycourobilin to C83 within MpeA in blue light | ^14^ |
| slr0151 | Involve in PSII assembly and repair (tetratricopeptide repeat protein on thylakoid ultrastructure during PS II assembly and repair) | ^16^ |
| CyanoP | Involve in the Early Steps of Photosystem II Assembly in the Cyanobacterium | ^17^ |

**Table S 7** Novel photosynthetic protein prediction using different methods with varying thresholds

| **Method** | **Threshold** | **Accuracy** | **F1 (minor class)** | **MCC** |
| --- | --- | --- | --- | --- |
| Blastp | E-value <= 100 | 0.902 | 0.000 | 0.001 |
|  | **E-value <= 10** | **0.789** | **0.188** | **0.078** |
|  | E-value <= 1 | 0.423 | 0.124 | -0.096 |
|  | E-value <= 0.1 | 0.252 | 0.080 | -0.279 |
|  | E-value <= 0.01 | 0.228 | 0.078 | -0.305 |
|  | E-value <= 0.001 | 0.228 | 0.078 | -0.305 |
| SVMProt* | **Probability >= 50%** | **0.650** | **0.218** | **0.104** |
|  | Probability >= 60% | 0.846 | 0.174 | 0.089 |
|  | Probability >= 70% | 0.829 | 0.087 | -0.007 |
|  | Probability >= 80% | 0.772 | 0.067 | -0.059 |
|  | Probability >= 90% | 0.675 | 0.000 | -0.179 |
| PHOTOMOD | E-value <= 10 | 0.789 | 0.235 | 0.133 |
|  | **E-value <= 1** | **0.772** | **0.300** | **0.214** |
|  | E-value <= 0.1 | 0.748 | 0.279 | 0.188 |
|  | E-value <= 0.01 | 0.659 | 0.222 | 0.110 |
|  | E-value <= 0.001 | 0.220 | 0.094 | -0.269 |
| *True positive is counted if predicted GO terms of positive input contain at least one of 61 photosynthetic GO terms, whereas false positive is counted if at least one of 61 photosynthetic GO terms is found in predicted GO terms of negative input. | | | | |

**Table S 8** The result of four methods in the prediction of novel photosynthetic proteins

| **No** | **Protein** | **Actual class** | **Blast** | **SVMProt** | **SCMPSP** | **PHOTOMOD** |
| --- | --- | --- | --- | --- | --- | --- |
| 1 | apcB3 | photo | photo | photo | photo | photo |
| 2 | apcD4 | photo | photo | non_photo | non_photo | photo |
| 3 | CyanoP | photo | non_photo | photo | photo | non_photo |
| 4 | DpxA | photo | non_photo | non_photo | non_photo | non_photo |
| 5 | fciA | photo | non_photo | photo | non_photo | photo |
| 6 | fciB | photo | non_photo | non_photo | non_photo | photo |
| 7 | IflA | photo | non_photo | non_photo | non_photo | non_photo |
| 8 | isiX | photo | non_photo | photo | photo | photo |
| 9 | MpeZ | photo | non_photo | photo | non_photo | photo |
| 10 | rfpA | photo | non_photo | non_photo | non_photo | non_photo |
| 11 | rfpB | photo | photo | photo | non_photo | non_photo |
| 12 | slr0151 | photo | non_photo | non_photo | non_photo | non_photo |
| 13 | A0A0D4BS77 | non_photo | non_photo | non_photo | non_photo | non_photo |
| 14 | A0A0D4BSN8 | non_photo | non_photo | non_photo | non_photo | non_photo |
| 15 | A0A0H3AJC2 | non_photo | non_photo | non_photo | non_photo | non_photo |
| 16 | A0A0H3AKU6 | non_photo | non_photo | non_photo | photo | non_photo |
| 17 | A0A0S3QTC6 | non_photo | non_photo | photo | non_photo | non_photo |
| 18 | A0A0S3QTD0 | non_photo | non_photo | non_photo | non_photo | photo |
| 19 | A0A1E7MYN1 | non_photo | non_photo | photo | non_photo | non_photo |
| 20 | A0QTU7 | non_photo | non_photo | photo | non_photo | non_photo |
| 21 | A0QTV0 | non_photo | non_photo | photo | photo | non_photo |
| 22 | A0R3Y2 | non_photo | non_photo | non_photo | non_photo | non_photo |
| 23 | A7AZH2 | non_photo | non_photo | non_photo | non_photo | non_photo |
| 24 | A7B3K3 | non_photo | non_photo | non_photo | non_photo | non_photo |
| 25 | A7NH01 | non_photo | non_photo | non_photo | non_photo | photo |
| 26 | A9AWD5 | non_photo | non_photo | non_photo | non_photo | non_photo |
| 27 | A9AWD6 | non_photo | non_photo | non_photo | non_photo | NotPredict |
| 28 | A9AWD7 | non_photo | photo | non_photo | non_photo | non_photo |
| 29 | A9FZ87 | non_photo | non_photo | non_photo | non_photo | photo |
| 30 | B1VXR4 | non_photo | non_photo | non_photo | non_photo | non_photo |
| 31 | B1W3T1 | non_photo | photo | non_photo | non_photo | non_photo |
| 32 | B2FI29 | non_photo | non_photo | non_photo | photo | non_photo |
| 33 | B5H7H3 | non_photo | photo | non_photo | photo | photo |
| 34 | B5HDJ6 | non_photo | non_photo | non_photo | non_photo | photo |
| 35 | B7J3C9 | non_photo | non_photo | photo | photo | non_photo |
| 36 | C7PLV2 | non_photo | non_photo | non_photo | photo | photo |
| 37 | C8WGQ3 | non_photo | non_photo | photo | non_photo | non_photo |
| 38 | C8WJW0 | non_photo | non_photo | photo | non_photo | non_photo |
| 39 | C8WMP0 | non_photo | non_photo | non_photo | non_photo | non_photo |
| 40 | D2B747 | non_photo | photo | photo | photo | photo |
| 41 | D2PPM7 | non_photo | non_photo | non_photo | non_photo | non_photo |
| 42 | D2PPM8 | non_photo | non_photo | photo | non_photo | photo |
| 43 | D8GR66 | non_photo | non_photo | photo | non_photo | non_photo |
| 44 | D8GR67 | non_photo | photo | photo | photo | non_photo |
| 45 | D8GR68 | non_photo | non_photo | non_photo | photo | non_photo |
| 46 | D8GR69 | non_photo | photo | non_photo | photo | non_photo |
| 47 | D8GR70 | non_photo | non_photo | non_photo | photo | non_photo |
| 48 | D8GR71 | non_photo | photo | non_photo | non_photo | non_photo |
| 49 | D9XD61 | non_photo | non_photo | non_photo | non_photo | photo |
| 50 | D9XDR8 | non_photo | non_photo | non_photo | non_photo | photo |
| 51 | E4N7E5 | non_photo | non_photo | photo | non_photo | photo |
| 52 | E8W6C7 | non_photo | non_photo | photo | non_photo | photo |
| 53 | E9RFT0 | non_photo | non_photo | non_photo | non_photo | non_photo |
| 54 | F9US27 | non_photo | non_photo | photo | non_photo | non_photo |
| 55 | F9UT67 | non_photo | non_photo | non_photo | non_photo | photo |
| 56 | F9UT68 | non_photo | non_photo | photo | non_photo | non_photo |
| 57 | H2A7G5 | non_photo | non_photo | non_photo | non_photo | NotPredict |
| 58 | H2K885 | non_photo | non_photo | non_photo | non_photo | non_photo |
| 59 | H2K888 | non_photo | non_photo | non_photo | non_photo | non_photo |
| 60 | H6LC27 | non_photo | non_photo | photo | non_photo | non_photo |
| 61 | H6LC29 | non_photo | non_photo | NotPredict | photo | non_photo |
| 62 | H6LC30 | non_photo | photo | non_photo | photo | non_photo |
| 63 | H6LC31 | non_photo | photo | non_photo | photo | non_photo |
| 64 | H6LC32 | non_photo | non_photo | non_photo | non_photo | non_photo |
| 65 | K0K750 | non_photo | non_photo | non_photo | non_photo | photo |
| 66 | K4JY29 | non_photo | photo | NotPredict | photo | NotPredict |
| 67 | K4REQ6 | non_photo | non_photo | non_photo | photo | photo |
| 68 | L8EUQ6 | non_photo | non_photo | photo | non_photo | non_photo |
| 69 | L8EYU3 | non_photo | non_photo | non_photo | non_photo | non_photo |
| 70 | O34138 | non_photo | non_photo | non_photo | non_photo | non_photo |
| 71 | O51767 | non_photo | non_photo | non_photo | non_photo | non_photo |
| 72 | O54143 | non_photo | non_photo | non_photo | photo | non_photo |
| 73 | O83323 | non_photo | non_photo | non_photo | photo | non_photo |
| 74 | O83324 | non_photo | non_photo | non_photo | photo | non_photo |
| 75 | P0DPE4 | non_photo | non_photo | photo | non_photo | non_photo |
| 76 | P0DPE9 | non_photo | photo | photo | photo | non_photo |
| 77 | P0DPF0 | non_photo | non_photo | non_photo | non_photo | non_photo |
| 78 | P71889 | non_photo | non_photo | non_photo | non_photo | non_photo |
| 79 | P96072 | non_photo | non_photo | photo | non_photo | non_photo |
| 80 | Q0P9Y2 | non_photo | non_photo | non_photo | photo | photo |
| 81 | Q0SJK9 | non_photo | non_photo | non_photo | non_photo | non_photo |
| 82 | Q0ZQ46 | non_photo | non_photo | non_photo | non_photo | non_photo |
| 83 | Q15JF5 | non_photo | non_photo | photo | photo | non_photo |
| 84 | Q15JF8 | non_photo | non_photo | photo | non_photo | non_photo |
| 85 | Q2PWU9 | non_photo | non_photo | non_photo | non_photo | non_photo |
| 86 | Q31KC7 | non_photo | non_photo | NotPredict | non_photo | photo |
| 87 | Q3K999 | non_photo | non_photo | NotPredict | non_photo | non_photo |
| 88 | Q3T6E2 | non_photo | non_photo | non_photo | non_photo | non_photo |
| 89 | Q46085 | non_photo | non_photo | NotPredict | non_photo | non_photo |
| 90 | Q4KCY6 | non_photo | non_photo | photo | non_photo | non_photo |
| 91 | Q4VKU9 | non_photo | non_photo | non_photo | non_photo | non_photo |
| 92 | Q4VKV0 | non_photo | non_photo | photo | non_photo | non_photo |
| 93 | Q4VKV1 | non_photo | photo | photo | photo | non_photo |
| 94 | Q5SHW0 | non_photo | non_photo | non_photo | photo | non_photo |
| 95 | Q5SIP0 | non_photo | non_photo | non_photo | non_photo | non_photo |
| 96 | Q65YW9 | non_photo | non_photo | photo | non_photo | non_photo |
| 97 | Q65YX0 | non_photo | non_photo | non_photo | photo | non_photo |
| 98 | Q6EZC2 | non_photo | non_photo | non_photo | non_photo | non_photo |
| 99 | Q6EZC3 | non_photo | non_photo | non_photo | photo | non_photo |
| 100 | Q72EF3 | non_photo | non_photo | photo | non_photo | photo |
| 101 | Q72EF4 | non_photo | non_photo | non_photo | non_photo | non_photo |
| 102 | Q768S8 | non_photo | non_photo | non_photo | non_photo | non_photo |
| 103 | Q768T3 | non_photo | non_photo | non_photo | non_photo | non_photo |
| 104 | Q7N561 | non_photo | non_photo | non_photo | non_photo | photo |
| 105 | Q81L64 | non_photo | photo | non_photo | photo | non_photo |
| 106 | Q81L65 | non_photo | non_photo | photo | non_photo | non_photo |
| 107 | Q81LM1 | non_photo | non_photo | non_photo | non_photo | non_photo |
| 108 | Q81QL7 | non_photo | non_photo | non_photo | photo | non_photo |
| 109 | Q81XB1 | non_photo | non_photo | non_photo | photo | non_photo |
| 110 | Q81XB2 | non_photo | NotPredict | non_photo | photo | non_photo |
| 111 | Q81XB3 | non_photo | non_photo | photo | non_photo | non_photo |
| 112 | Q826W3 | non_photo | non_photo | non_photo | non_photo | non_photo |
| 113 | Q845S8 | non_photo | non_photo | non_photo | photo | non_photo |
| 114 | Q845S9 | non_photo | non_photo | non_photo | non_photo | non_photo |
| 115 | Q899Y1 | non_photo | non_photo | photo | photo | non_photo |
| 116 | Q8A712 | non_photo | non_photo | non_photo | non_photo | non_photo |
| 117 | Q8GHB1 | non_photo | non_photo | non_photo | non_photo | non_photo |
| 118 | Q8PHA1 | non_photo | non_photo | non_photo | non_photo | non_photo |
| 119 | Q9AF95 | non_photo | photo | non_photo | photo | non_photo |
| 120 | Q9AGW3 | non_photo | photo | photo | photo | non_photo |
| 121 | Q9LCB4 | non_photo | non_photo | non_photo | non_photo | non_photo |
| 122 | Q9X721 | non_photo | photo | NotPredict | photo | non_photo |
| 123 | S4S3E3 | non_photo | non_photo | non_photo | non_photo | non_photo |
| *Green color is used to emphasize the correct prediction, while red color is used to emphasize incorrect prediction. | | | | | | |

**Table S 9** Prediction of unknown proteins in cyanobacteria genome, *Synechocystis* sp. PCC 6803

| **Query** | **totalBlastHits** | **PhotoHits(%)** | **nonphotoHits(%)** | **noPredictHits** | **prediction** | **Ave_Prob_prediction** |
| --- | --- | --- | --- | --- | --- | --- |
| sgl0002 | 33 | 57.576 | 42.424 | 0 | photo | 0.683 |
| sll0037 | 143 | 52.448 | 47.552 | 0 | photo | 0.666 |
| sll0069 | 78 | 96.154 | 3.846 | 0 | photo | 0.698 |
| sll0088 | 138 | 78.261 | 21.739 | 0 | photo | 0.941 |
| sll0149 | 60 | 96.667 | 3.333 | 0 | photo | 0.562 |
| sll0160 | 64 | 92.187 | 7.812 | 0 | photo | 0.806 |
| sll0178 | 24 | 54.167 | 45.833 | 0 | photo | 0.682 |
| sll0185 | 126 | 100.000 | 0.000 | 0 | photo | 0.961 |
| sll0253 | 36 | 97.222 | 2.778 | 0 | photo | 0.663 |
| sll0272 | 94 | 100.000 | 0.000 | 0 | photo | 0.822 |
| sll0318 | 146 | 100.000 | 0.000 | 0 | photo | 0.734 |
| sll0364 | 111 | 67.568 | 32.432 | 0 | photo | 0.760 |
| sll0413 | 136 | 93.382 | 6.618 | 0 | photo | 0.553 |
| sll0423 | 48 | 93.750 | 6.250 | 0 | photo | 0.672 |
| sll0436 | 66 | 100.000 | 0.000 | 0 | photo | 0.915 |
| sll0442 | 45 | 93.333 | 6.667 | 0 | photo | 0.673 |
| sll0471 | 162 | 87.037 | 12.963 | 0 | photo | 0.680 |
| sll0497 | 91 | 97.802 | 2.198 | 0 | photo | 0.658 |
| sll0528 | 263 | 55.133 | 44.867 | 0 | photo | 0.671 |
| sll0543 | 18 | 88.889 | 11.111 | 0 | photo | 0.750 |
| sll0544 | 121 | 80.992 | 19.008 | 0 | photo | 0.587 |
| sll0572 | 14 | 92.857 | 7.143 | 0 | photo | 0.669 |
| sll0585 | 107 | 86.916 | 13.084 | 0 | photo | 0.778 |
| sll0611 | 60 | 98.333 | 1.667 | 0 | photo | 0.897 |
| sll0615 | 208 | 69.712 | 30.288 | 0 | photo | 0.852 |
| sll0639 | 56 | 92.857 | 7.143 | 0 | photo | 0.670 |
| sll0696 | 160 | 56.250 | 43.750 | 0 | photo | 0.639 |
| sll0732 | 17 | 88.235 | 11.765 | 0 | photo | 0.673 |
| sll0814 | 73 | 100.000 | 0.000 | 0 | photo | 0.741 |
| sll0853 | 203 | 60.591 | 39.409 | 0 | photo | 0.795 |
| sll0860 | 98 | 71.429 | 28.571 | 0 | photo | 0.665 |
| sll0861 | 109 | 88.991 | 11.009 | 0 | photo | 0.574 |
| sll0997 | 38 | 94.737 | 5.263 | 0 | photo | 0.554 |
| sll1060 | 90 | 98.889 | 1.111 | 0 | photo | 0.777 |
| sll1158 | 27 | 100.000 | 0.000 | 0 | photo | 0.654 |
| sll1242 | 558 | 68.638 | 31.362 | 0 | photo | 0.641 |
| sll1340 | 93 | 98.925 | 1.075 | 0 | photo | 0.940 |
| sll1372 | 98 | 69.388 | 30.612 | 0 | photo | 0.558 |
| sll1381 | 43 | 86.047 | 13.953 | 0 | photo | 0.550 |
| sll1389 | 24 | 100.000 | 0.000 | 0 | photo | 0.700 |
| sll1390 | 96 | 82.292 | 17.708 | 0 | photo | 0.683 |
| sll1399 | 114 | 100.000 | 0.000 | 0 | photo | 0.842 |
| sll1400 | 106 | 69.811 | 30.189 | 0 | photo | 0.817 |
| sll1414 | 94 | 100.000 | 0.000 | 0 | photo | 0.987 |
| sll1455 | 84 | 100.000 | 0.000 | 0 | photo | 0.654 |
| sll1485 | 129 | 99.225 | 0.775 | 0 | photo | 0.730 |
| sll1486 | 127 | 99.213 | 0.787 | 0 | photo | 0.732 |
| sll1500 | 95 | 98.947 | 1.053 | 0 | photo | 0.584 |
| sll1532 | 57 | 100.000 | 0.000 | 0 | photo | 0.654 |
| sll1543 | 52 | 94.231 | 5.769 | 0 | photo | 0.658 |
| sll1573 | 56 | 94.643 | 5.357 | 0 | photo | 0.847 |
| sll1608 | 106 | 88.679 | 11.321 | 0 | photo | 0.538 |
| sll1738 | 192 | 58.333 | 41.667 | 0 | photo | 0.735 |
| sll1757 | 95 | 96.842 | 3.158 | 0 | photo | 0.828 |
| sll1874 | 190 | 99.474 | 0.526 | 0 | photo | 0.934 |
| sll1913 | 279 | 55.556 | 44.444 | 0 | photo | 0.742 |
| sll1915 | 76 | 89.474 | 10.526 | 0 | photo | 0.742 |
| sll1916 | 137 | 64.964 | 35.036 | 0 | photo | 0.867 |
| sll1940 | 107 | 73.832 | 26.168 | 0 | photo | 0.712 |
| sll1979 | 97 | 92.784 | 7.216 | 0 | photo | 0.869 |
| sll5003 | 124 | 71.774 | 28.226 | 0 | photo | 0.812 |
| sll5030 | 10 | 100.000 | 0.000 | 0 | photo | 0.654 |
| sll5032 | 98 | 86.735 | 13.265 | 0 | photo | 0.613 |
| sll5033 | 80 | 100.000 | 0.000 | 0 | photo | 0.701 |
| sll5034 | 49 | 55.102 | 44.898 | 0 | photo | 0.676 |
| sll5097 | 89 | 91.011 | 8.989 | 0 | photo | 0.664 |
| sll5130 | 6 | 100.000 | 0.000 | 0 | photo | 0.654 |
| sll5132 | 54 | 96.296 | 3.704 | 0 | photo | 0.654 |
| sll6054 | 7 | 57.143 | 42.857 | 0 | photo | 0.659 |
| sll6055 | 10 | 70.000 | 30.000 | 0 | photo | 0.738 |
| sll7031 | 59 | 98.305 | 1.695 | 0 | photo | 0.656 |
| sll7033 | 112 | 60.714 | 39.286 | 0 | photo | 0.754 |
| sll7069 | 15 | 100.000 | 0.000 | 0 | photo | 0.654 |
| sll8004 | 39 | 69.231 | 30.769 | 0 | photo | 0.696 |
| sll8018 | 4 | 100.000 | 0.000 | 0 | photo | 0.654 |
| sll8019 | 5 | 80.000 | 20.000 | 0 | photo | 0.717 |
| sll8025 | 22 | 95.455 | 4.545 | 0 | photo | 0.650 |
| sll8032 | 5 | 100.000 | 0.000 | 0 | photo | 0.654 |
| sll8035 | 8 | 100.000 | 0.000 | 0 | photo | 0.654 |
| slr0022 | 94 | 97.872 | 2.128 | 0 | photo | 0.867 |
| slr0076 | 290 | 99.655 | 0.345 | 0 | photo | 0.988 |
| slr0142 | 37 | 51.351 | 48.649 | 0 | photo | 0.686 |
| slr0144 | 112 | 98.214 | 1.786 | 0 | photo | 0.768 |
| slr0146 | 67 | 98.507 | 1.493 | 0 | photo | 0.710 |
| slr0147 | 93 | 96.774 | 3.226 | 0 | photo | 0.745 |
| slr0148 | 293 | 63.823 | 36.177 | 0 | photo | 0.741 |
| slr0149 | 250 | 97.600 | 2.400 | 0 | photo | 0.914 |
| slr0211 | 89 | 77.528 | 22.472 | 0 | photo | 0.653 |
| slr0249 | 95 | 100.000 | 0.000 | 0 | photo | 0.821 |
| slr0263 | 54 | 98.148 | 1.852 | 0 | photo | 0.616 |
| slr0299 | 8 | 100.000 | 0.000 | 0 | photo | 0.563 |
| slr0320 | 606 | 57.591 | 42.409 | 0 | photo | 0.703 |
| slr0356 | 139 | 74.101 | 25.899 | 0 | photo | 0.763 |
| slr0363 | 93 | 95.699 | 4.301 | 0 | photo | 0.660 |
| slr0388 | 138 | 99.275 | 0.725 | 0 | photo | 0.858 |
| slr0397 | 62 | 100.000 | 0.000 | 0 | photo | 0.544 |
| slr0404 | 107 | 73.832 | 26.168 | 0 | photo | 0.553 |
| slr0438 | 89 | 100.000 | 0.000 | 0 | photo | 0.553 |
| slr0554 | 99 | 93.939 | 6.061 | 0 | photo | 0.665 |
| slr0598 | 94 | 98.936 | 1.064 | 0 | photo | 0.864 |
| slr0619 | 65 | 60.000 | 40.000 | 0 | photo | 0.769 |
| slr0625 | 43 | 97.674 | 2.326 | 0 | photo | 0.679 |
| slr0642 | 78 | 88.462 | 11.538 | 0 | photo | 0.687 |
| slr0725 | 28 | 100.000 | 0.000 | 0 | photo | 0.654 |
| slr0732 | 90 | 100.000 | 0.000 | 0 | photo | 0.651 |
| slr0770 | 392 | 50.255 | 49.745 | 0 | photo | 0.830 |
| slr0771 | 157 | 67.516 | 32.484 | 0 | photo | 0.735 |
| slr0780 | 136 | 58.824 | 41.176 | 0 | photo | 0.656 |
| slr0863 | 457 | 94.967 | 5.033 | 0 | photo | 0.664 |
| slr0869 | 76 | 78.947 | 21.053 | 0 | photo | 0.818 |
| slr0975 | 65 | 100.000 | 0.000 | 0 | photo | 0.654 |
| slr1100 | 32 | 100.000 | 0.000 | 0 | photo | 0.893 |
| slr1170 | 241 | 71.078 | 28.922 | 37 | photo | 0.854 |
| slr1173 | 53 | 94.340 | 5.660 | 0 | photo | 0.656 |
| slr1174 | 95 | 87.368 | 12.632 | 0 | photo | 0.655 |
| slr1182 | 115 | 75.652 | 24.348 | 0 | photo | 0.702 |
| slr1186 | 50 | 98.000 | 2.000 | 0 | photo | 0.808 |
| slr1188 | 165 | 97.576 | 2.424 | 0 | photo | 0.741 |
| slr1195 | 95 | 96.842 | 3.158 | 0 | photo | 0.861 |
| slr1220 | 116 | 81.034 | 18.966 | 0 | photo | 0.850 |
| slr1260 | 80 | 100.000 | 0.000 | 0 | photo | 0.701 |
| slr1261 | 501 | 51.209 | 48.791 | 46 | photo | 0.691 |
| slr1266 | 125 | 84.000 | 16.000 | 0 | photo | 0.754 |
| slr1353 | 120 | 91.667 | 8.333 | 0 | photo | 0.659 |
| slr1394 | 94 | 98.936 | 1.064 | 0 | photo | 0.861 |
| slr1413 | 128 | 78.906 | 21.094 | 0 | photo | 0.725 |
| slr1451 | 69 | 89.855 | 10.145 | 0 | photo | 0.782 |
| slr1478 | 49 | 91.837 | 8.163 | 0 | photo | 0.662 |
| slr1495 | 107 | 85.981 | 14.019 | 0 | photo | 0.948 |
| slr1546 | 79 | 92.405 | 7.595 | 0 | photo | 0.633 |
| slr1570 | 100 | 68.000 | 32.000 | 0 | photo | 0.817 |
| slr1572 | 95 | 71.579 | 28.421 | 0 | photo | 0.815 |
| slr1601 | 71 | 95.775 | 4.225 | 0 | photo | 0.947 |
| slr1624 | 82 | 82.927 | 17.073 | 0 | photo | 0.672 |
| slr1676 | 140 | 60.714 | 39.286 | 0 | photo | 0.638 |
| slr1699 | 118 | 98.305 | 1.695 | 0 | photo | 0.911 |
| slr1732 | 69 | 100.000 | 0.000 | 0 | photo | 0.845 |
| slr1767 | 104 | 71.154 | 28.846 | 0 | photo | 0.844 |
| slr1770 | 63 | 96.825 | 3.175 | 0 | photo | 0.654 |
| slr1799 | 149 | 75.168 | 24.832 | 0 | photo | 0.594 |
| slr1800 | 84 | 97.619 | 2.381 | 0 | photo | 0.769 |
| slr1880 | 54 | 90.741 | 9.259 | 0 | photo | 0.663 |
| slr1885 | 50 | 98.000 | 2.000 | 0 | photo | 0.579 |
| slr1907 | 57 | 100.000 | 0.000 | 0 | photo | 0.654 |
| slr1923 | 114 | 94.737 | 5.263 | 0 | photo | 0.517 |
| slr1927 | 130 | 54.615 | 45.385 | 0 | photo | 0.745 |
| slr1949 | 144 | 99.306 | 0.694 | 0 | photo | 0.741 |
| slr1998 | 391 | 65.473 | 34.527 | 0 | photo | 0.796 |
| slr2000 | 110 | 67.273 | 32.727 | 0 | photo | 0.662 |
| slr2003 | 37 | 100.000 | 0.000 | 0 | photo | 0.645 |
| slr2025 | 64 | 73.437 | 26.562 | 0 | photo | 0.524 |
| slr2052 | 44 | 97.727 | 2.273 | 0 | photo | 0.655 |
| slr2070 | 51 | 86.275 | 13.725 | 0 | photo | 0.815 |
| slr5012 | 16 | 100.000 | 0.000 | 0 | photo | 0.654 |
| slr5087 | 9 | 77.778 | 22.222 | 0 | photo | 0.689 |
| slr5101 | 11 | 100.000 | 0.000 | 0 | photo | 0.654 |
| slr5102 | 12 | 100.000 | 0.000 | 0 | photo | 0.654 |
| slr6029 | 6 | 83.333 | 16.667 | 0 | photo | 0.699 |
| slr6049 | 22 | 59.091 | 40.909 | 0 | photo | 0.647 |
| slr6051 | 3 | 100.000 | 0.000 | 0 | photo | 0.654 |
| slr6057 | 9 | 66.667 | 33.333 | 0 | photo | 0.700 |
| slr6088 | 6 | 83.333 | 16.667 | 0 | photo | 0.699 |
| slr6094 | 41 | 53.659 | 46.341 | 0 | photo | 0.722 |
| slr6104 | 7 | 85.714 | 14.286 | 0 | photo | 0.664 |
| slr6106 | 32 | 100.000 | 0.000 | 0 | photo | 0.654 |
| slr7012 | 36 | 100.000 | 0.000 | 0 | photo | 0.654 |
| slr7013 | 34 | 100.000 | 0.000 | 0 | photo | 0.654 |
| slr7023 | 34 | 76.471 | 23.529 | 0 | photo | 0.669 |
| slr7024 | 65 | 87.692 | 12.308 | 0 | photo | 0.644 |
| slr7037 | 93 | 95.699 | 4.301 | 0 | photo | 0.661 |
| slr7058 | 35 | 54.286 | 45.714 | 0 | photo | 0.727 |
| slr7059 | 15 | 66.667 | 33.333 | 0 | photo | 0.719 |
| slr7076 | 3 | 100.000 | 0.000 | 0 | photo | 0.669 |
| slr7091 | 16 | 93.750 | 6.250 | 0 | photo | 0.659 |
| slr7094 | 10 | 100.000 | 0.000 | 0 | photo | 0.654 |
| slr7096 | 12 | 100.000 | 0.000 | 0 | photo | 0.690 |
| slr7097 | 33 | 100.000 | 0.000 | 0 | photo | 0.654 |
| smr0015 | 32 | 84.375 | 15.625 | 0 | photo | 0.720 |
| ssl0241 | 88 | 72.727 | 27.273 | 0 | photo | 0.687 |
| ssl0294 | 60 | 95.000 | 5.000 | 0 | photo | 0.660 |
| ssl0312 | 53 | 100.000 | 0.000 | 0 | photo | 0.654 |
| ssl0385 | 110 | 77.273 | 22.727 | 0 | photo | 0.714 |
| ssl0483 | 49 | 67.347 | 32.653 | 0 | photo | 0.836 |
| ssl0511 | 30 | 96.667 | 3.333 | 0 | photo | 0.664 |
| ssl0788 | 69 | 94.203 | 5.797 | 0 | photo | 0.570 |
| ssl0900 | 11 | 81.818 | 18.182 | 0 | photo | 0.674 |
| ssl1004 | 147 | 87.075 | 12.925 | 0 | photo | 0.660 |
| ssl1046 | 13 | 92.308 | 7.692 | 0 | photo | 0.656 |
| ssl1376 | 177 | 50.847 | 49.153 | 0 | photo | 0.770 |
| ssl1378 | 64 | 95.312 | 4.687 | 0 | photo | 0.557 |
| ssl2009 | 94 | 79.787 | 20.213 | 0 | photo | 0.722 |
| ssl2648 | 81 | 51.852 | 48.148 | 0 | photo | 0.763 |
| ssl2717 | 66 | 95.455 | 4.545 | 0 | photo | 0.925 |
| ssl2920 | 70 | 100.000 | 0.000 | 0 | photo | 0.940 |
| ssl2921 | 130 | 71.538 | 28.462 | 0 | photo | 0.815 |
| ssl2999 | 24 | 95.833 | 4.167 | 0 | photo | 0.663 |
| ssl3297 | 26 | 100.000 | 0.000 | 0 | photo | 0.507 |
| ssl3719 | 126 | 50.794 | 49.206 | 0 | photo | 0.711 |
| ssl3803 | 26 | 84.615 | 15.385 | 0 | photo | 0.647 |
| ssl5031 | 85 | 97.647 | 2.353 | 0 | photo | 0.614 |
| ssl5095 | 33 | 96.970 | 3.030 | 0 | photo | 0.658 |
| ssl5099 | 38 | 55.263 | 44.737 | 0 | photo | 0.710 |
| ssl7042 | 2 | 100.000 | 0.000 | 0 | photo | 0.654 |
| ssl7048 | 15 | 100.000 | 0.000 | 0 | photo | 0.654 |
| ssl7053 | 4 | 100.000 | 0.000 | 0 | photo | 0.899 |
| ssl8028 | 16 | 100.000 | 0.000 | 0 | photo | 0.654 |
| ssr0755 | 66 | 57.576 | 42.424 | 0 | photo | 0.708 |
| ssr1558 | 237 | 75.527 | 24.473 | 0 | photo | 0.858 |
| ssr1698 | 109 | 77.982 | 22.018 | 0 | photo | 0.866 |
| ssr1880 | 61 | 95.082 | 4.918 | 0 | photo | 0.782 |
| ssr1951 | 59 | 100.000 | 0.000 | 0 | photo | 0.637 |
| ssr2009 | 77 | 50.649 | 49.351 | 0 | photo | 0.769 |
| ssr2062 | 71 | 94.366 | 5.634 | 0 | photo | 0.671 |
| ssr2781 | 64 | 100.000 | 0.000 | 0 | photo | 0.905 |
| ssr2806 | 69 | 79.710 | 20.290 | 0 | photo | 0.677 |
| ssr3189 | 87 | 63.218 | 36.782 | 0 | photo | 0.718 |
| ssr3304 | 255 | 66.667 | 33.333 | 0 | photo | 0.905 |
| ssr5011 | 81 | 100.000 | 0.000 | 0 | photo | 0.925 |
| ssr5019 | 249 | 99.197 | 0.803 | 0 | photo | 0.561 |
| ssr5106 | 15 | 86.667 | 13.333 | 0 | photo | 0.686 |
| ssr6046 | 15 | 86.667 | 13.333 | 0 | photo | 0.639 |
| ssr8013 | 35 | 68.571 | 31.429 | 0 | photo | 0.664 |
| slr1667 | 12 | 91.667 | 8.333 | 0 | photo | 0.678 |
| slr9003 | 88 | 95.455 | 4.545 | 0 | photo | 0.654 |
| ssr9004 | 5 | 100.000 | 0.000 | 0 | photo | 0.654 |
| sll0047 | 69 | 100.000 | 0.000 | 0 | photo | 0.883 |
| sll1509 | 109 | 99.083 | 0.917 | 0 | photo | 0.889 |
| ssl1417 | 92 | 98.913 | 1.087 | 0 | photo | 0.548 |
| sll0584 | 91 | 82.418 | 17.582 | 0 | photo | 0.636 |
| slr0480 | 856 | 55.023 | 44.977 | 0 | photo | 0.755 |
| sll1214 | 189 | 100.000 | 0.000 | 0 | photo | 0.935 |
| slr0923 | 94 | 97.872 | 2.128 | 0 | photo | 0.659 |
| sll0101 | 35 | 74.286 | 25.714 | 0 | photo | 0.675 |
| sll0167 | 11 | 100.000 | 0.000 | 0 | photo | 0.579 |
| sll0225 | 33 | 93.939 | 6.061 | 0 | photo | 0.884 |
| sll0263 | 21 | 85.714 | 14.286 | 0 | photo | 0.911 |
| sll0265 | 22 | 90.909 | 9.091 | 0 | photo | 0.879 |
| sll0266 | 20 | 100.000 | 0.000 | 0 | photo | 0.903 |
| sll0280 | 54 | 87.037 | 12.963 | 0 | photo | 0.657 |
| sll0327 | 25 | 60.000 | 40.000 | 0 | photo | 0.734 |
| sll0328 | 15 | 93.333 | 6.667 | 0 | photo | 0.835 |
| sll0403 | 8 | 100.000 | 0.000 | 0 | photo | 0.654 |
| sll0405 | 9 | 88.889 | 11.111 | 0 | photo | 0.688 |
| sll0406 | 17 | 64.706 | 35.294 | 0 | photo | 0.669 |
| sll0419 | 7 | 100.000 | 0.000 | 0 | photo | 0.654 |
| sll0426 | 10 | 100.000 | 0.000 | 0 | photo | 0.654 |
| sll0443 | 13 | 100.000 | 0.000 | 0 | photo | 0.654 |
| sll0444 | 9 | 100.000 | 0.000 | 0 | photo | 0.654 |
| sll0445 | 22 | 68.182 | 31.818 | 0 | photo | 0.710 |
| sll0446 | 23 | 78.261 | 21.739 | 0 | photo | 0.707 |
| sll0447 | 19 | 94.737 | 5.263 | 0 | photo | 0.659 |
| sll0448 | 20 | 95.000 | 5.000 | 0 | photo | 0.657 |
| sll0449 | 16 | 100.000 | 0.000 | 0 | photo | 0.654 |
| sll0494 | 8 | 100.000 | 0.000 | 0 | photo | 0.654 |
| sll0508 | 9 | 100.000 | 0.000 | 0 | photo | 0.654 |
| sll0539 | 11 | 81.818 | 18.182 | 0 | photo | 0.837 |
| sll0552 | 10 | 90.000 | 10.000 | 0 | photo | 0.764 |
| sll0563 | 17 | 100.000 | 0.000 | 0 | photo | 0.654 |
| sll0588 | 14 | 100.000 | 0.000 | 0 | photo | 0.678 |
| sll0595 | 41 | 51.220 | 48.780 | 0 | photo | 0.685 |
| sll0614 | 18 | 100.000 | 0.000 | 0 | photo | 0.584 |
| sll0623 | 56 | 100.000 | 0.000 | 0 | photo | 0.864 |
| sll0625 | 18 | 83.333 | 16.667 | 0 | photo | 0.674 |
| sll0630 | 12 | 100.000 | 0.000 | 0 | photo | 0.654 |
| sll0722 | 9 | 88.889 | 11.111 | 0 | photo | 0.683 |
| sll0733 | 36 | 97.222 | 2.778 | 0 | photo | 0.657 |
| sll0847 | 95 | 67.368 | 32.632 | 0 | photo | 0.668 |
| sll0872 | 18 | 55.556 | 44.444 | 0 | photo | 0.756 |
| sll0943 | 16 | 93.750 | 6.250 | 0 | photo | 0.659 |
| sll0980 | 28 | 82.143 | 17.857 | 0 | photo | 0.659 |
| sll0981 | 12 | 75.000 | 25.000 | 0 | photo | 0.698 |
| sll1061 | 21 | 90.476 | 9.524 | 0 | photo | 0.663 |
| sll1062 | 19 | 100.000 | 0.000 | 0 | photo | 0.654 |
| sll1086 | 60 | 100.000 | 0.000 | 0 | photo | 0.654 |
| sll1132 | 15 | 100.000 | 0.000 | 0 | photo | 0.654 |
| sll1163 | 40 | 57.500 | 42.500 | 0 | photo | 0.706 |
| sll1239 | 15 | 53.333 | 46.667 | 0 | photo | 0.781 |
| sll1240 | 8 | 100.000 | 0.000 | 0 | photo | 0.654 |
| sll1241 | 9 | 88.889 | 11.111 | 0 | photo | 0.679 |
| sll1267 | 13 | 92.308 | 7.692 | 0 | photo | 0.670 |
| sll1268 | 74 | 95.946 | 4.054 | 0 | photo | 0.948 |
| sll1272 | 11 | 100.000 | 0.000 | 0 | photo | 0.654 |
| sll1273 | 20 | 90.000 | 10.000 | 0 | photo | 0.664 |
| sll1338 | 23 | 100.000 | 0.000 | 0 | photo | 0.654 |
| sll1359 | 122 | 72.131 | 27.869 | 0 | photo | 0.699 |
| sll1373 | 8 | 100.000 | 0.000 | 0 | photo | 0.534 |
| sll1396 | 12 | 83.333 | 16.667 | 0 | photo | 0.711 |
| sll1401 | 8 | 87.500 | 12.500 | 0 | photo | 0.923 |
| sll1476 | 1 | 100.000 | 0.000 | 0 | photo | 0.654 |
| sll1531 | 26 | 96.154 | 3.846 | 0 | photo | 0.664 |
| sll1665 | 22 | 81.818 | 18.182 | 0 | photo | 0.671 |
| sll1714 | 26 | 100.000 | 0.000 | 0 | photo | 0.654 |
| sll1717 | 23 | 73.913 | 26.087 | 0 | photo | 0.661 |
| sll1730 | 28 | 89.286 | 10.714 | 0 | photo | 0.671 |
| sll1755 | 17 | 100.000 | 0.000 | 0 | photo | 0.654 |
| sll1761 | 11 | 100.000 | 0.000 | 0 | photo | 0.654 |
| sll1763 | 35 | 82.857 | 17.143 | 0 | photo | 0.745 |
| sll1764 | 36 | 97.222 | 2.778 | 0 | photo | 0.798 |
| sll1765 | 8 | 100.000 | 0.000 | 0 | photo | 0.906 |
| sll1830 | 53 | 98.113 | 1.887 | 0 | photo | 0.817 |
| sll1853 | 19 | 100.000 | 0.000 | 0 | photo | 0.666 |
| sll1863 | 51 | 96.078 | 3.922 | 0 | photo | 0.659 |
| sll1885 | 35 | 100.000 | 0.000 | 0 | photo | 0.654 |
| sll5002 | 11 | 81.818 | 18.182 | 0 | photo | 0.692 |
| sll5006 | 10 | 80.000 | 20.000 | 0 | photo | 0.697 |
| sll5044 | 9 | 55.556 | 44.444 | 0 | photo | 0.732 |
| sll5069 | 3 | 66.667 | 33.333 | 0 | photo | 0.638 |
| sll5089 | 20 | 95.000 | 5.000 | 0 | photo | 0.660 |
| sll5090 | 7 | 100.000 | 0.000 | 0 | photo | 0.654 |
| sll5109 | 3 | 66.667 | 33.333 | 0 | photo | 0.638 |
| sll6010 | 7 | 100.000 | 0.000 | 0 | photo | 0.654 |
| sll6069 | 7 | 100.000 | 0.000 | 0 | photo | 0.654 |
| sll7009 | 23 | 100.000 | 0.000 | 0 | photo | 0.654 |
| sll7043 | 30 | 86.667 | 13.333 | 0 | photo | 0.769 |
| sll7062 | 18 | 83.333 | 16.667 | 0 | photo | 0.674 |
| sll7063 | 37 | 100.000 | 0.000 | 0 | photo | 0.654 |
| sll7064 | 6 | 100.000 | 0.000 | 0 | photo | 0.654 |
| sll7066 | 41 | 75.610 | 24.390 | 0 | photo | 0.712 |
| sll7067 | 24 | 100.000 | 0.000 | 0 | photo | 0.654 |
| sll7070 | 26 | 96.154 | 3.846 | 0 | photo | 0.860 |
| sll7085 | 25 | 96.000 | 4.000 | 0 | photo | 0.663 |
| sll7086 | 10 | 100.000 | 0.000 | 0 | photo | 0.654 |
| sll7087 | 37 | 94.595 | 5.405 | 0 | photo | 0.658 |
| sll7089 | 26 | 96.154 | 3.846 | 0 | photo | 0.660 |
| sll7090 | 31 | 100.000 | 0.000 | 0 | photo | 0.654 |
| sll8007 | 22 | 90.909 | 9.091 | 0 | photo | 0.681 |
| sll8011 | 45 | 91.111 | 8.889 | 0 | photo | 0.716 |
| sll8017 | 1 | 100.000 | 0.000 | 0 | photo | 0.654 |
| sll8033 | 18 | 100.000 | 0.000 | 0 | photo | 0.654 |
| slr0019 | 18 | 77.778 | 22.222 | 0 | photo | 0.700 |
| slr0059 | 28 | 100.000 | 0.000 | 0 | photo | 0.717 |
| slr0061 | 17 | 100.000 | 0.000 | 0 | photo | 0.654 |
| slr0103 | 11 | 72.727 | 27.273 | 0 | photo | 0.673 |
| slr0145 | 26 | 100.000 | 0.000 | 0 | photo | 0.884 |
| slr0196 | 23 | 100.000 | 0.000 | 0 | photo | 0.828 |
| slr0209 | 66 | 96.970 | 3.030 | 0 | photo | 0.655 |
| slr0226 | 80 | 97.500 | 2.500 | 0 | photo | 0.945 |
| slr0262 | 24 | 87.500 | 12.500 | 0 | photo | 0.617 |
| slr0271 | 9 | 100.000 | 0.000 | 0 | photo | 0.654 |
| slr0272 | 10 | 80.000 | 20.000 | 0 | photo | 0.662 |
| slr0386 | 66 | 71.212 | 28.788 | 0 | photo | 0.650 |
| slr0398 | 16 | 87.500 | 12.500 | 0 | photo | 0.649 |
| slr0421 | 67 | 64.179 | 35.821 | 0 | photo | 0.758 |
| slr0442 | 136 | 62.500 | 37.500 | 0 | photo | 0.821 |
| slr0496 | 15 | 73.333 | 26.667 | 0 | photo | 0.715 |
| slr0522 | 22 | 86.364 | 13.636 | 0 | photo | 0.676 |
| slr0569 | 21 | 80.952 | 19.048 | 0 | photo | 0.681 |
| slr0572 | 10 | 100.000 | 0.000 | 0 | photo | 0.654 |
| slr0573 | 8 | 100.000 | 0.000 | 0 | photo | 0.654 |
| slr0579 | 8 | 100.000 | 0.000 | 0 | photo | 0.654 |
| slr0581 | 15 | 86.667 | 13.333 | 0 | photo | 0.676 |
| slr0582 | 8 | 100.000 | 0.000 | 0 | photo | 0.654 |
| slr0602 | 10 | 80.000 | 20.000 | 0 | photo | 0.924 |
| slr0667 | 9 | 100.000 | 0.000 | 0 | photo | 0.654 |
| slr0668 | 8 | 100.000 | 0.000 | 0 | photo | 0.654 |
| slr0727 | 10 | 70.000 | 30.000 | 0 | photo | 0.693 |
| slr0881 | 89 | 76.404 | 23.596 | 0 | photo | 0.638 |
| slr0912 | 21 | 66.667 | 33.333 | 0 | photo | 0.729 |
| slr0913 | 8 | 87.500 | 12.500 | 0 | photo | 0.689 |
| slr0914 | 12 | 91.667 | 8.333 | 0 | photo | 0.668 |
| slr0937 | 53 | 71.698 | 28.302 | 0 | photo | 0.658 |
| slr1107 | 15 | 100.000 | 0.000 | 0 | photo | 0.663 |
| slr1135 | 19 | 73.684 | 26.316 | 0 | photo | 0.725 |
| slr1168 | 113 | 93.805 | 6.195 | 0 | photo | 0.801 |
| slr1187 | 9 | 100.000 | 0.000 | 0 | photo | 0.785 |
| slr1210 | 84 | 53.571 | 46.429 | 0 | photo | 0.767 |
| slr1222 | 9 | 100.000 | 0.000 | 0 | photo | 0.654 |
| slr1232 | 10 | 90.000 | 10.000 | 0 | photo | 0.681 |
| slr1243 | 90 | 53.333 | 46.667 | 0 | photo | 0.656 |
| slr1396 | 13 | 84.615 | 15.385 | 0 | photo | 0.640 |
| slr1397 | 8 | 100.000 | 0.000 | 0 | photo | 0.654 |
| slr1398 | 7 | 100.000 | 0.000 | 0 | photo | 0.654 |
| slr1421 | 35 | 100.000 | 0.000 | 0 | photo | 0.685 |
| slr1437 | 43 | 100.000 | 0.000 | 42 | photo | 0.654 |
| slr1450 | 19 | 100.000 | 0.000 | 0 | photo | 0.654 |
| slr1544 | 35 | 97.143 | 2.857 | 0 | photo | 0.658 |
| slr1552 | 26 | 100.000 | 0.000 | 0 | photo | 0.654 |
| slr1567 | 37 | 94.595 | 5.405 | 0 | photo | 0.893 |
| slr1571 | 84 | 79.762 | 20.238 | 0 | photo | 0.811 |
| slr1576 | 9 | 88.889 | 11.111 | 0 | photo | 0.807 |
| slr1616 | 119 | 85.714 | 14.286 | 0 | photo | 0.608 |
| slr1670 | 36 | 83.333 | 16.667 | 0 | photo | 0.660 |
| slr1773 | 32 | 100.000 | 0.000 | 0 | photo | 0.654 |
| slr1774 | 36 | 77.778 | 22.222 | 0 | photo | 0.672 |
| slr1778 | 12 | 100.000 | 0.000 | 0 | photo | 0.548 |
| slr1788 | 65 | 98.462 | 1.538 | 0 | photo | 0.657 |
| slr1789 | 65 | 98.462 | 1.538 | 0 | photo | 0.657 |
| slr1869 | 9 | 88.889 | 11.111 | 0 | photo | 0.673 |
| slr1920 | 10 | 100.000 | 0.000 | 0 | photo | 0.663 |
| slr1959 | 56 | 64.286 | 35.714 | 0 | photo | 0.717 |
| slr2018 | 39 | 100.000 | 0.000 | 0 | photo | 0.823 |
| slr2037 | 21 | 71.429 | 28.571 | 0 | photo | 0.658 |
| slr5013 | 21 | 80.952 | 19.048 | 0 | photo | 0.634 |
| slr5016 | 46 | 86.957 | 13.043 | 0 | photo | 0.677 |
| slr5073 | 11 | 100.000 | 0.000 | 0 | photo | 0.641 |
| slr5085 | 3 | 100.000 | 0.000 | 0 | photo | 0.654 |
| slr5126 | 2 | 100.000 | 0.000 | 0 | photo | 0.654 |
| slr6004 | 1 | 100.000 | 0.000 | 0 | photo | 0.654 |
| slr6005 | 13 | 92.308 | 7.692 | 0 | photo | 0.677 |
| slr6006 | 14 | 92.857 | 7.143 | 0 | photo | 0.676 |
| slr6007 | 49 | 63.265 | 36.735 | 0 | photo | 0.780 |
| slr6013 | 6 | 83.333 | 16.667 | 0 | photo | 0.702 |
| slr6014 | 10 | 70.000 | 30.000 | 0 | photo | 0.729 |
| slr6015 | 7 | 100.000 | 0.000 | 0 | photo | 0.654 |
| slr6016 | 11 | 100.000 | 0.000 | 0 | photo | 0.654 |
| slr6021 | 8 | 100.000 | 0.000 | 0 | photo | 0.654 |
| slr6022 | 4 | 100.000 | 0.000 | 0 | photo | 0.654 |
| slr6028 | 10 | 80.000 | 20.000 | 0 | photo | 0.676 |
| slr6031 | 17 | 76.471 | 23.529 | 0 | photo | 0.706 |
| slr6033 | 1 | 100.000 | 0.000 | 0 | photo | 0.654 |
| slr6045 | 3 | 100.000 | 0.000 | 0 | photo | 0.711 |
| slr6063 | 9 | 88.889 | 11.111 | 0 | photo | 0.685 |
| slr6064 | 13 | 92.308 | 7.692 | 0 | photo | 0.677 |
| slr6065 | 14 | 92.857 | 7.143 | 0 | photo | 0.676 |
| slr6066 | 49 | 63.265 | 36.735 | 0 | photo | 0.780 |
| slr6072 | 6 | 83.333 | 16.667 | 0 | photo | 0.702 |
| slr6073 | 10 | 70.000 | 30.000 | 0 | photo | 0.729 |
| slr6074 | 7 | 100.000 | 0.000 | 0 | photo | 0.654 |
| slr6075 | 11 | 100.000 | 0.000 | 0 | photo | 0.654 |
| slr6080 | 8 | 100.000 | 0.000 | 0 | photo | 0.654 |
| slr6081 | 4 | 100.000 | 0.000 | 0 | photo | 0.654 |
| slr6087 | 10 | 80.000 | 20.000 | 0 | photo | 0.676 |
| slr6090 | 15 | 86.667 | 13.333 | 0 | photo | 0.684 |
| slr6091 | 2 | 100.000 | 0.000 | 0 | photo | 0.654 |
| slr7010 | 103 | 59.223 | 40.777 | 0 | photo | 0.727 |
| slr7011 | 30 | 96.667 | 3.333 | 0 | photo | 0.659 |
| slr7061 | 15 | 93.333 | 6.667 | 0 | photo | 0.676 |
| slr7080 | 28 | 100.000 | 0.000 | 0 | photo | 0.654 |
| slr7081 | 8 | 62.500 | 37.500 | 0 | photo | 0.667 |
| slr7099 | 2 | 100.000 | 0.000 | 0 | photo | 0.751 |
| slr7100 | 10 | 90.000 | 10.000 | 0 | photo | 0.688 |
| ssl0467 | 31 | 87.097 | 12.903 | 0 | photo | 0.675 |
| ssl0750 | 8 | 100.000 | 0.000 | 0 | photo | 0.654 |
| ssl0787 | 76 | 88.158 | 11.842 | 0 | photo | 0.576 |
| ssl1326 | 18 | 55.556 | 44.444 | 0 | photo | 0.742 |
| ssl1533 | 26 | 92.308 | 7.692 | 0 | photo | 0.656 |
| ssl2138 | 66 | 96.970 | 3.030 | 0 | photo | 0.758 |
| ssl2245 | 34 | 97.059 | 2.941 | 0 | photo | 0.663 |
| ssl2420 | 55 | 87.273 | 12.727 | 0 | photo | 0.692 |
| ssl2501 | 25 | 100.000 | 0.000 | 0 | photo | 0.654 |
| ssl2507 | 37 | 100.000 | 0.000 | 0 | photo | 0.654 |
| ssl3142 | 10 | 100.000 | 0.000 | 0 | photo | 0.821 |
| ssl3222 | 15 | 93.333 | 6.667 | 0 | photo | 0.772 |
| ssl3383 | 15 | 86.667 | 13.333 | 0 | photo | 0.673 |
| ssl3615 | 18 | 72.222 | 27.778 | 0 | photo | 0.645 |
| ssl3769 | 25 | 100.000 | 0.000 | 0 | photo | 0.654 |
| ssl5001 | 20 | 100.000 | 0.000 | 0 | photo | 0.513 |
| ssl5007 | 1 | 100.000 | 0.000 | 0 | photo | 0.654 |
| ssl5065 | 6 | 83.333 | 16.667 | 0 | photo | 0.683 |
| ssl5091 | 1 | 100.000 | 0.000 | 0 | photo | 0.654 |
| ssl5096 | 32 | 87.500 | 12.500 | 0 | photo | 0.691 |
| ssl5098 | 11 | 90.909 | 9.091 | 0 | photo | 0.646 |
| ssl5103 | 15 | 93.333 | 6.667 | 0 | photo | 0.666 |
| ssl6023 | 17 | 88.235 | 11.765 | 0 | photo | 0.660 |
| ssl6035 | 11 | 100.000 | 0.000 | 0 | photo | 0.654 |
| ssl6082 | 17 | 88.235 | 11.765 | 0 | photo | 0.660 |
| ssl6092 | 20 | 55.000 | 45.000 | 0 | photo | 0.732 |
| ssr0693 | 11 | 90.909 | 9.091 | 0 | photo | 0.651 |
| ssr1038 | 25 | 96.000 | 4.000 | 0 | photo | 0.666 |
| ssr1049 | 36 | 69.444 | 30.556 | 0 | photo | 0.695 |
| ssr1853 | 26 | 76.923 | 23.077 | 0 | photo | 0.692 |
| ssr2049 | 255 | 65.098 | 34.902 | 0 | photo | 0.902 |
| ssr2153 | 11 | 100.000 | 0.000 | 0 | photo | 0.654 |
| ssr2194 | 32 | 100.000 | 0.000 | 0 | photo | 0.654 |
| ssr2317 | 51 | 80.392 | 19.608 | 0 | photo | 0.668 |
| ssr2406 | 23 | 100.000 | 0.000 | 0 | photo | 0.654 |
| ssr2422 | 50 | 78.000 | 22.000 | 0 | photo | 0.696 |
| ssr2975 | 12 | 100.000 | 0.000 | 0 | photo | 0.573 |
| ssr3159 | 8 | 87.500 | 12.500 | 0 | photo | 0.667 |
| ssr3465 | 8 | 100.000 | 0.000 | 0 | photo | 0.654 |
| ssr3532 | 54 | 88.889 | 11.111 | 0 | photo | 0.653 |
| ssr5074 | 3 | 100.000 | 0.000 | 0 | photo | 0.654 |
| ssr6002 | 2 | 100.000 | 0.000 | 0 | photo | 0.654 |
| ssr6003 | 10 | 70.000 | 30.000 | 0 | photo | 0.747 |
| ssr6048 | 9 | 77.778 | 22.222 | 0 | photo | 0.693 |
| ssr7035 | 1 | 100.000 | 0.000 | 0 | photo | 0.654 |
| slr9203 | 5 | 60.000 | 40.000 | 0 | photo | 0.768 |
| slr9102 | 5 | 80.000 | 20.000 | 0 | photo | 0.696 |
| slr9002 | 1 | 100.000 | 0.000 | 0 | photo | 0.654 |
